# The structural basis of RanGAP1 regulation and catalysis in nuclear transport

**DOI:** 10.64898/2026.04.28.721414

**Authors:** Liang Xu, Hyunbum Jang, Ruth Nussinov

## Abstract

RanGAP1 promotes GTP hydrolysis of nuclear pore complex (NPC) transport complexes at the cytoplasmic face. A disordered linker connects its catalytic GAP domain to the C-terminal sumoylation domain, anchoring into NPC’s cytoplasmic filaments. This arrangement raises the question of how these distinct functions are coordinated within a crowded cellular environment. Using atomistic molecular dynamics simulations, we show that RanGAP1 adopts an autoinhibited conformation, where the C-terminal domain masks the catalytic GAP domain. Sumoylation allosterically relieves this autoinhibition, enabling GTP-bound Ran access to the GAP domain. In the cytosol, Ran-GTP/RanBP1 can bind a less populated open conformation of RanGAP1, providing a backup mechanism for GTP hydrolysis in Ran. Importantly, we observe that Arg191 of human RanGAP1 inserts into the GTP-binding pocket of Ran and directly interacts with the γ-phosphate, consistent with a canonical arginine finger. This observation contrasts with earlier models derived from yeast RanGAP and suggests that human RanGAP1 may follow a catalytic mechanism similar to classical small GTPase regulators like NF1. Together, these findings provide a framework of RanGAP1, linking autoinhibition, sumoylation, spatial organization at the NPC, and the catalytic mechanism. They also highlight how conformational regulation and post-translational modification coordinate efficient GTP hydrolysis in Ran during nuclear transport.

## Introduction

The structures of human RanGAP1 (Ran GTPase-activating protein 1) and RasGAP (Ras GTPase-activating protein NF1) are distinct. Studies of yeast-based predominantly cytosolic RanGAP (Rna1p) suggested that their catalytic mechanisms also differ. Unlike NF1, which uses a canonical arginine finger to activate Ras, yeast RanGAP uses an invariant glutamine (Gln69) within its leucine-rich repeat (LRR), a mechanism previously assumed to apply to human RanGAP1. Understanding this mechanism is crucial, as Ran serves as the primary regulator of nucleocytoplasmic transport through the nuclear pore complex (NPC). Our extensive molecular dynamics simulations, supported by available experimental data, lead us to propose that the human RanGAP1 catalytic mechanism uses the canonical arginine (Arg191) finger, similar to its functional analog, NF1. The highly conserved Gln69 can contribute to GTP hydrolysis by orienting the nucleophilic water molecule toward the γ-phosphate of GTP, like Gln61 of K-Ras4B. Here we reveal the structural basis of RanGAP1 regulation and catalysis in nuclear transport.

Nucleocytoplasmic transport is an essential and highly regulated process that maintains cellular homeostasis by controlling the exchange of proteins and RNA between the nucleus and cytoplasm^1–4^. The transport is mediated by NPC^2,5–7^, a large protein assembly embedded in the nuclear envelope. The NPC forms a central transport channel flanked by cytoplasmic filaments that captures incoming transport complexes, and a nuclear basket, a filamentous assembly on the nuclear side that facilitates cargo release and transport^8,9^. Directional cargo transport is driven by the Ran GTPase cycle and nuclear transport receptors^10,11^. The guanine nucleotide exchange factor (GEF) for Ran, regulator of chromosome condensation 1 (RCC1), is localized in the nucleus, where it activates Ran by catalyzing the exchange of GDP for GTP. RanGAP1 stimulates the hydrolysis of GTP in the cytoplasm^12–14^. Thus, RCC1 and RanGAP1 together establish the asymmetric distribution of Ran-GTP in the nucleus and Ran-GDP in the cytoplasm, providing directionality of cargo transport^15^. Recent work has uncovered a noncanonical function of RanGAP1 in forming a Ras-GTP/RanGAP1 complex that enhances nuclear export, underscoring its broader role in regulating nucleocytoplasmic transport and cellular signaling^16^.

A critical structural feature on the cytoplasmic face of the NPC is the cytoplasmic filament^17^, primarily composed of Ran binding protein 2 (RanBP2, also known as Nup358). In human cells, RanGAP1 undergoes small ubiquitin-like modifier 1 (SUMO1) modification at its C-terminal domain that is essential for its stable association with the E2 enzyme Ubc9 and RanBP2 at the NPC^18–24^. Importantly, RanBP2 contains a SUMO E3 ligase domain that promotes SUMO1 conjugation of RanGAP1. The SUMO-conjugated RanGAP1/SUMO1/Ubc9/RanBP2 assembly functions as a SUMO E3 ligase^25–27^, as well as a disassembly machine for export complexes (**Fig. 1A**). RanGAP1 stimulates GTP hydrolysis in Ran, dismantling the CRM1/Ran-GTP/cargo complex^28^. Thus, RanBP2 coordinates the spatial organization of Ran-GTP binding, SUMO1 conjugation, and export complex disassembly on the cytoplasmic side of the NPC, a process in which RanGAP1 plays an essential role.

**Fig. 1.**
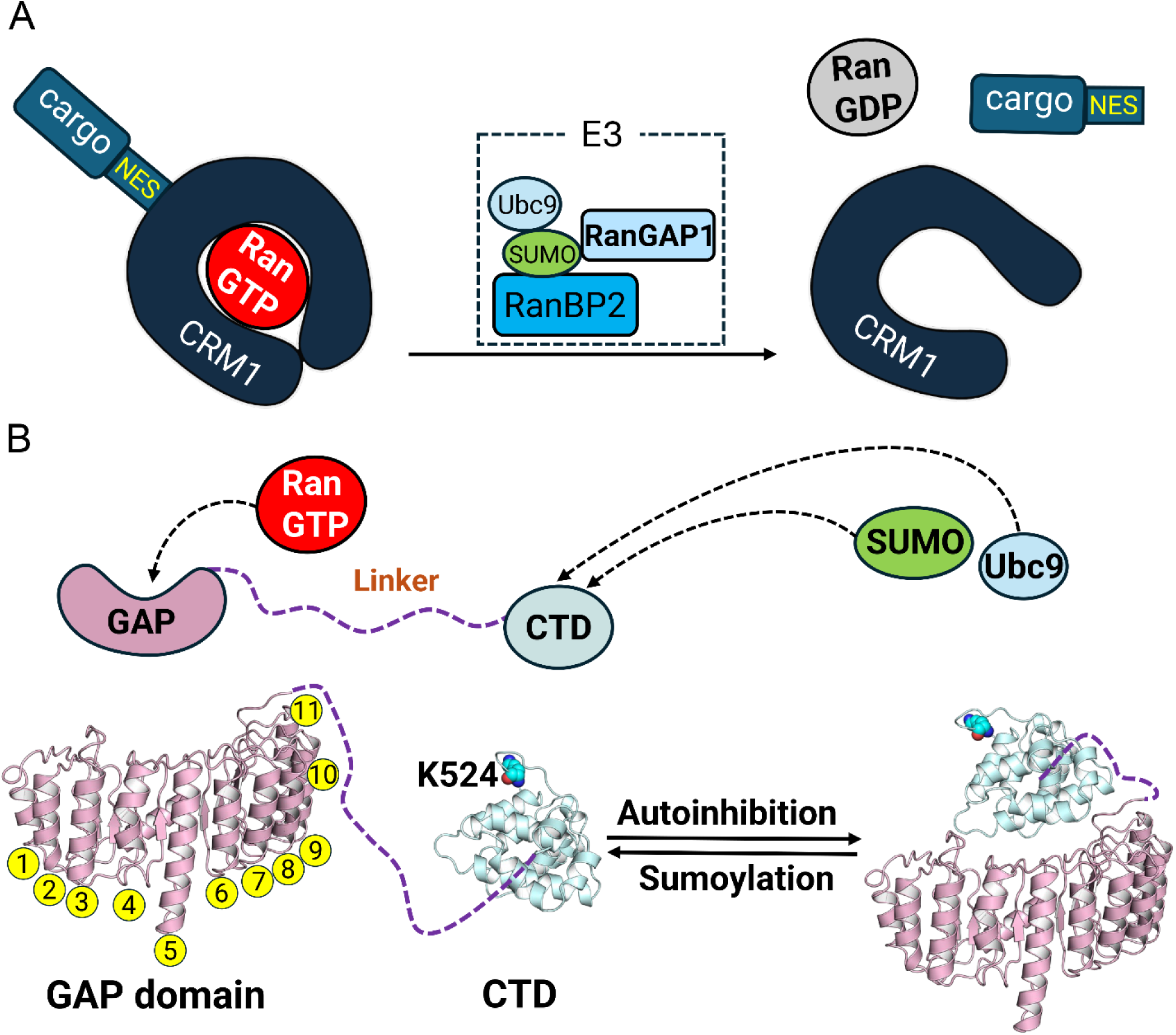
RanGAP1 is essential for nucleocytoplasmic transport and can adopt an autoinhibited conformation in a crowded cellular environment to function more efficiently. (**A**) Dual functional roles of the SUMO-conjugated RanGAP1/SUMO1/Ubc9/RanBP2 complex. This complex acts as a SUMO E3 ligase promoting SUMO1 conjugation, and a platform that facilitates disassembly of the CRM1/Ran-GTP/cargo export complex. CRM1 (export receptor chromosomal region maintenance 1, also known as XPO1, Exportin 1) is the export receptor. The NES (nuclear export signal) is the structural motif in the cargo that is recognized by CRM1. (**B**) Domain organization of human RanGAP1. RanGAP1 comprises an N-terminal GAP domain (residues 1–362), a disordered linker region, and a C-terminal sumoylation domain (CTD, residues 432–587) that mediates its localization to the NPC. The GAP domain consists of eleven leucine-rich repeats. The sumoylation site, Lys524, is in the C-terminal of RanGAP1. We propose that RanGAP adopts an autoinhibited conformation prior to sumoylation, which may contribute to its functional efficiency.

Current structural and functional insights into RanGAP were mostly obtained from studies of yeast^29–34^. However, yeast Rna1p only consists of the GAP domain and does not require sumoylation for its NPC architecture, which lacks RanBP2^33^. In contrast, human RanGAP1 is sumoylated at its C-terminal domain and attached to RanBP2, representing the main pathway for GTP hydrolysis in Ran in the cytoplasm^35,36^. Unmodified RanGAP1 also functions in the cytosol, where RanBP1 binds Ran-GTP and facilitates formation of a RanGAP1/Ran-GTP/RanBP1 complex, promoting GTP hydrolysis in Ran and potentially serving as a backup mechanism. Notably, RanGAP1 contains a long-disordered region between its N-terminal GAP domain and its C-terminal sumoylation domain (**Fig. 1B**). In the crowded condensate environment, maintaining coordination between these two functional modules is critical for efficient coupling between SUMO1-mediated attachment to RanBP2 and GTP hydrolysis. To achieve this, RanGAP1 could adopt a transient autoinhibited conformation in which its GAP domain interacts with its C-terminal region prior to sumoylation (**Fig. 1B**). Such an autoinhibition maintains proximity between the two domains but restraining RanGAP1 activity. Sumoylation can mediate protein-protein interactions,^37^ but it remains unclear whether sumoylation at the NPC could relieve RanGAP1 autoinhibition and allow RanGAP1 to associate with RanBP2 and hydrolyze GTP in Ran.

For many GTPases, such as Ras and Rho, the GAPs provide an essential arginine residue—termed the ‘arginine finger’—to accelerate the catalytic rate. While early studies of Rna1p suggested that GAP-mediated GTP hydrolysis in Ran does not require a classic arginine finger^30^, recent structural data on a human RanGAP1-containing complex present a more nuanced view^38^. This structure captures a state analogous to the ground OFF state of the GAP-related domain of the neurofibromin 1 (NF1), a Ras GAP^39^. In this conformation, the equivalent arginine residue, Arg191, does not contact the nucleotide^38^. Thus, it remains unresolved whether RanGAP1-mediated GTP hydrolysis utilizes a non-traditional arginine finger or follows the classic arginine finger-dependent mechanism.

Our all-atom molecular dynamics (MD) simulations reveal how the intramolecular interactions between the N-terminal GAP domain and the C-terminal sumoylation domain coordinate RanGAP1 function during nucleocytoplasmic transport. Sumoylation can allosterically relieve RanGAP1 autoinhibition, enabling the sumoylated C-terminal domain association with RanBP2 and the GAP domain interaction with Ran-GTP. We further determined the molecular mechanism underlying RanGAP1-mediated GTP hydrolysis at the cytoplasmic face of NPC, and establish a potential arginine finger involved in this process. This mechanism favors a catalytic mechanism similar to classical small GTPase regulators like NF1, differing from the yeast-based model.

## Results

### RanGAP1 can adopt various autoinhibited conformations prior to sumoylation

To elucidate the structural basis of RanGAP autoinhibition, we employed Rosetta docking to map the most favorable interactions between the GAP and C-terminal domains. From the resulting decoys, five top-ranked structural models (M1–M5) were selected (**Fig. 2A**). Across these models, the C-terminal domain is positioned at distinct regions of the GAP domain, suggesting multiple potential binding interfaces. In M1 to M3, the C-terminal domain primarily associates with LRRs 3–8 in the GAP domain, a site that overlaps with the binding region for Ran-GTP in complex with RanGAP1. In contrast, in M4 and M5, the C-terminal domain docks against the convex and concave surfaces of the GAP domain, respectively. We quantitatively evaluated these docking poses using two Rosetta docking scores: the total score (T_sc), representing the overall energetic stability of the entire complex, and the interface score (I_sc), which specifically measures the quality of the fit and interactions directly at the binding interface (**Fig. 2B**). Among these models, M1 seems the most favorable, showing the lowest T_sc (-1165) and I_sc (-32). As Rosetta docking scores represent a simplified energetic model to measure stability and binding affinity between the two domains of RanGAP1, we then performed all-atom MD simulations to further evaluate these models.

**Fig. 2.**
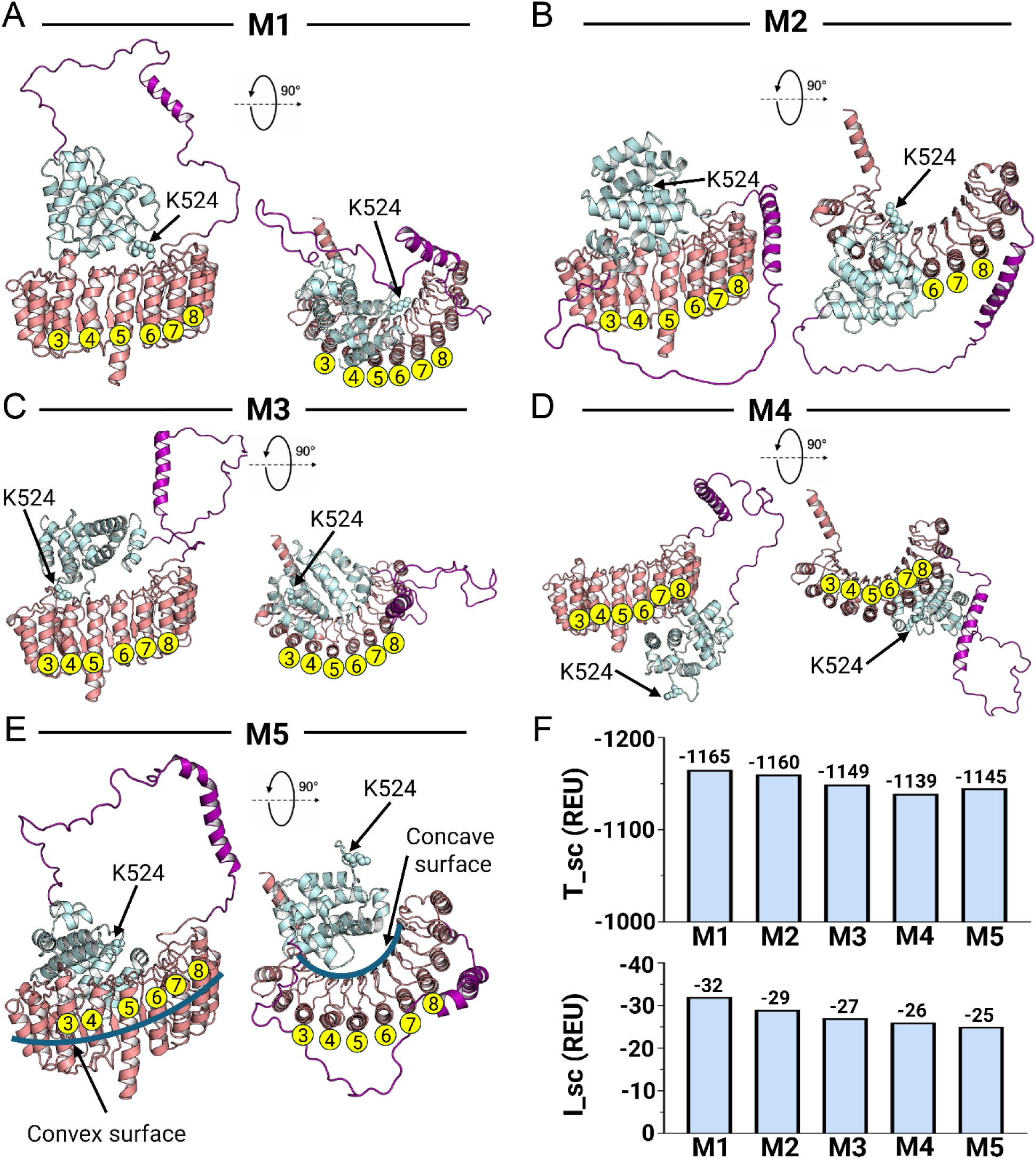
Structural models of autoinhibited RanGAP1. (**A**–**E**) The top five conformations of the autoinhibited RanGAP1 models (M1–M5) obtained from Rosetta docking, shown in side and top views. The sumoylation site Lys524 is labeled in each model. **(F)** Bar plots represent the Rosetta scores for M1–M5. The total score T_sc represents the overall structural stability of the model, while the interface score I_sc assesses the interactions between the GAP and the C-terminal domains of RanGAP1. Rosetta scores are reported in Rosetta Energy Unit (REU), which is a dimensionless, empirical energy scale derived from Rosetta scoring fuction. The leucine-rich repeats 3–8 were labeled to show the relative position of the C-terminal domain with respect to the GAP domain of RanGAP1.

We observed that M1, M3, M4, and M5 maintain stable autoinhibited conformations (**Figs. 3A–D**). For M2, however, the C-terminal domain quickly dissociated from the GAP domain during simulations, suggesting unfavorable interdomain interactions within this model (**Fig. 3E**). Since MD simulations allow full atomic flexibility and time-dependent relaxation under more physiologically relevant conditions than Rosetta docking, conformational changes in these autoinhibited RanGAP1 models are expected. We then performed clustering analysis on the trajectory of each model using the clustering algorithm by Daura et al.^40^, with the disordered linker region excluded. Conformations were grouped based on pairwise root-mean-square deviation (RMSD) using a defined cutoff (3 Å), and the most representative conformations of autoinhibited RanGAP1 were taken from the largest cluster. For each model (M1, M3, M4, and M5), we found only one dominant cluster representing the most populated conformational ensemble, indicating relatively stable conformations of autoinhibited RanGAP1. Compared to the initial structure, we observed substantial conformational rearrangement in M3 in which the C-terminal domain shifted its binding interface from the initial LRR 3–8 to LRR 6–10 (**Fig. 3B**). To improve sampling of the representative conformations of autoinhibited RanGAP1, we used these conformations as starting structures and performed three independent replicate simulations for M1, M3, M4, and M5, respectively. Among these models, M5 shows the largest buried surface area (3,348 Å²), higher than M1 (1,603 Å²), M3 (1,118 Å²), and M4 (1,942 Å²), suggesting a stronger association between the C-terminal and GAP domains (**Fig. S1**).

**Fig. 3.**
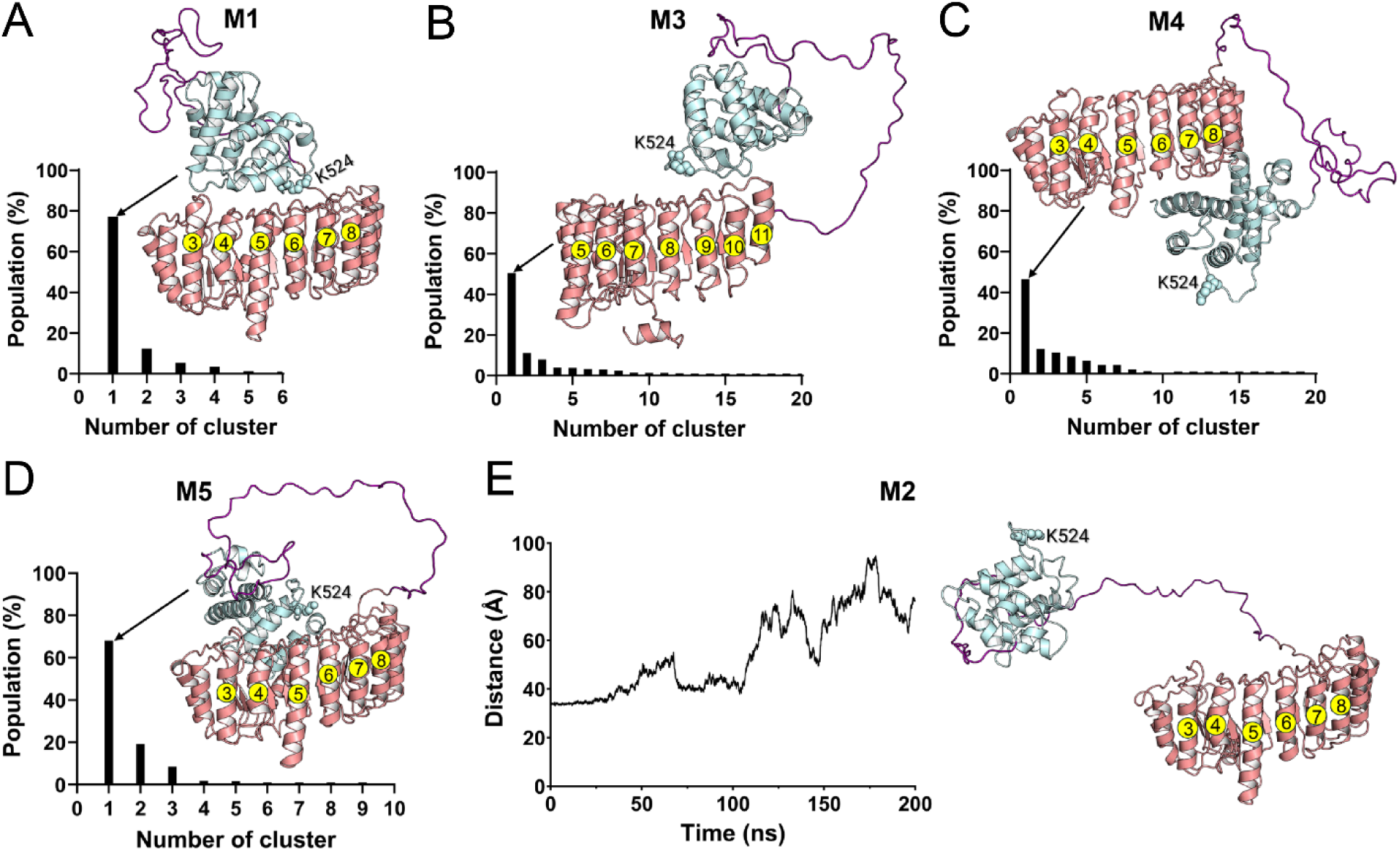
RanGAP1 adopts different autoinhibited conformations. Interactions between domains were found to be stable in M1 (**A**), M3 (**B**), M4 (**C**), and M5 (**D**). For M2 (**E**), the C-terminal domain dissociates from the GAP domain during MD simulations. Representative conformations are selected from the most populated (top 1) cluster for each system. Clustering analysis indicates that this dominant cluster is highly populated in all systems.

Besides the buried surface areas, we further characterized the autoinhibitory interface between the GAP and C-terminal domains by analyzing the populations of complementary electrostatic interactions and their contributions to the autoinhibited conformations of RanGAP1. The variations in the populations among three independent replicas reflect a dynamic autoinhibitory interface between the two domains with certain interactions persistently maintained. Despite distinct autoinhibitory interfaces observed in M1, M3, M4 and M5, the GAP domain interacts with the C-terminal domain via different loop regions linking the β-strand and α-helix within each LRR repeat. For M1, Asp245, Asp273 and Glu304 from the loops of LRRs 8–10, as well as Asp359 and Asp360 at the end of the GAP domain, interact with three lysine residues of the C-terminal domain (Lys524, Lys528, and Lys565), with Asp245^GAP^–Lys565^CTD^ and Asp273^GAP^–Lys565^CTD^ interactions most stable (**Fig. 4A**). In M3, more positively charged residues from the loops of LRRs 6–10 are involved in the interactions with the C-terminal domain, yet roughly half of these interactions occur with contact frequencies less than 50%, indicating transient autoinhibitory interactions. The most stable interactions occur between Asp245 and Lys528, as well as between Asp273 and Lys528 (**Fig. 4B**), a shift from Lys565 in M1. In contrast to M1 and M3, the C-terminal domain in M4 interacts with the GAP domain on the opposite side of the LRR (**Fig. 3C**), with loop residues on LRRs 6–10 interacting with the C-terminal domain (**Fig. 4C**). The autoinhibitory interface in M4 is stabilized by the interactions involving Arg264^GAP^–Asp495^CTD^, Arg264^GAP^–Asp492^CTD^, Lys294^GAP^–Asp549^CTD^, and Asp319^GAP^–Lys452^CTD^. The autoinhibitory interface in M5 seems more dynamic as evidenced by significant fluctuations in the contact frequencies, indicating more transient domain interactions that contribute to this autoinhibited RanGAP1 (**Fig. 4D**). Residues Arg189 and Arg191, located in the loop of LRR6, interact with Asp482 of the C-terminal domain with average contact frequencies of 50% and 30%, respectively. Meanwhile, Lys528 and Lys530 of the C-terminal domain contribute to the autoinhibitory interface through interactions with Asp359 and Asp360 at the end of the GAP domain.

**Fig. 4.**
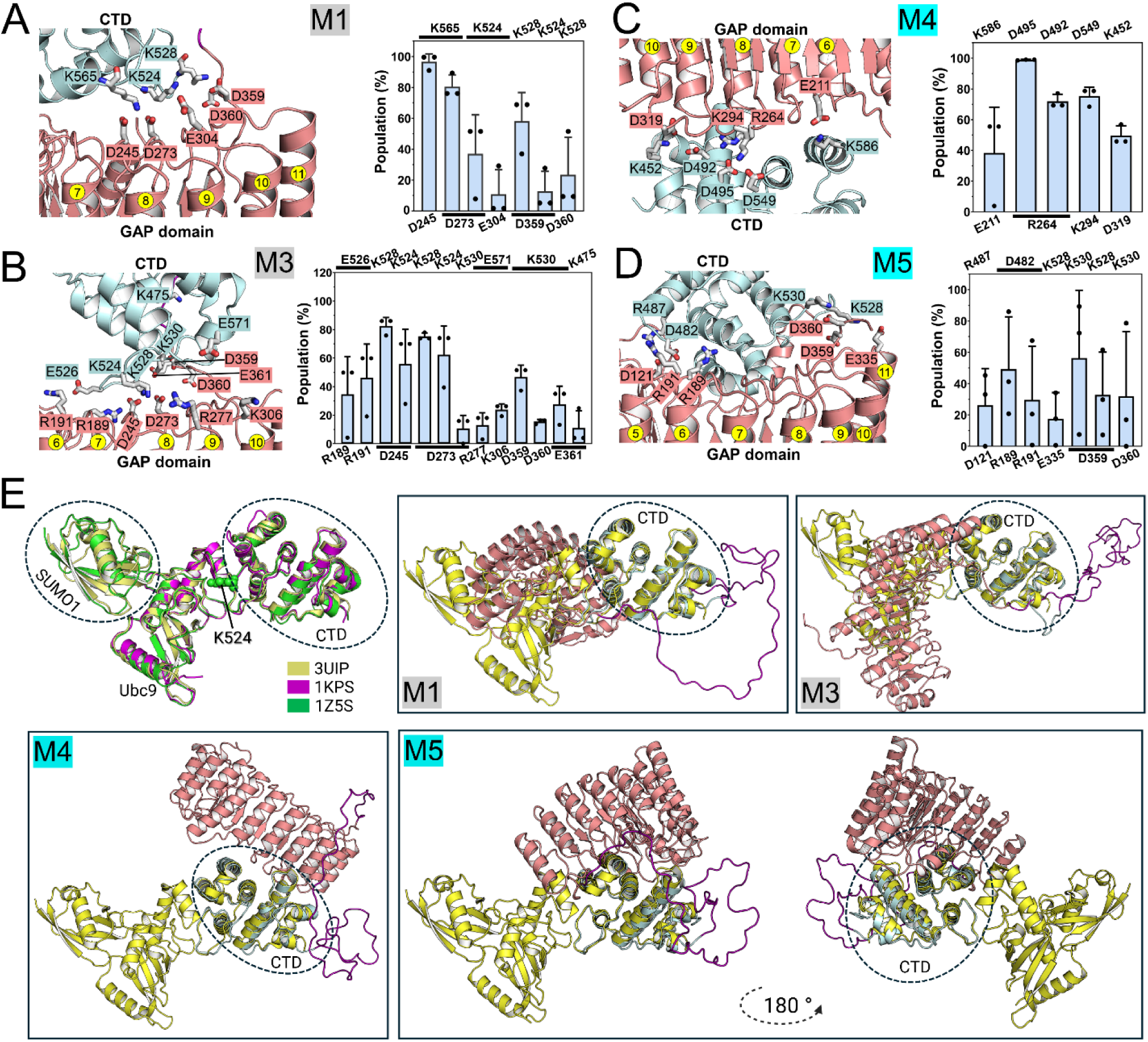
Autoinhibited RanGAP1 adopts sumoylation-competent conformation. (**A**–**D**) Complementary electrostatic interactions and their contributions to the autoinhibited conformations of RanGAP1 in M1 (**A**), M3 (**B**), M4 (**C**), and M5 (**D**). The population of each pair interaction is averaged over three independent replicate simulations for each system. (**E**) M4 and M5 adopt sumoylation-competent conformations. Structural studies indicate that Ubc9 forms a stable complex with SUMO1 and RanGAP1 C-terminal domain in a conserved spatial arrangement. Superposition of the three available crystal structures supports this conserved arrangement. Mapping this structural arrangement onto the autoinhibited RanGAP1 models shows that M1 and M3 are sumoylation-incompetent, whereas M4 and M5 adopt sumoylation-competent conformations.

Among the four autoinhibited models of RanGAP1, Lys524—the sumoylation site—is involved in the autoinhibitory interactions in M1 and M3 (**Figs. 4A** and **4B**), making it less accessible to SUMO1. In contrast, in M4 and M5 Lys524 does not participate in the domain interactions and remains accessible to SUMO1 (**Figs. 4C** and **4D**). These results imply that RanGAP1 could dynamically populate distinct autoinhibited states and modulate sumoylation site accessibility. To evaluate the structural feasibility of sumoylation in these models, we superimposed three crystal structures of sumoylated RanGAP1 bound to the E2 conjugating enzyme Ubc9 and/or SUMO1 (**Fig. 4E**), and found that they overlap quite well, indicating a highly conserved overall arrangement between RanGAP1, Ubc9, and SUMO1. In the crystal structure of the RanGAP1/Ubc9 complex (PDB: 1KPS)^41^, Ubc9 directly recognizes and binds the region surrounding Lys524 to position substrate lysine for SUMO1 transfer. Because Ubc9 remains associated with the C-terminal domain of RanGAP1 during SUMO1 transfer^42^, sumoylation requires formation of a ternary complex RanGAP1/SUMO1/Ubc9. Therefore, we superimposed this ternary complex with our proposed autoinhibited RanGAP1 models in terms of their C-terminal domains. As expected, in M1 and M3, the inaccessibility of Lys524 leads to significant steric clashes with the SUMO1/Ubc9 complex. In contrast, M4 and M5 show no steric clashes with the SUMO1/Ubc9 complex, suggesting that these conformations can readily accommodate the SUMO1/Ubc9 complex for SUMO1 conjugation. To elucidate how sumoylation modulates the autoinhibition of RanGAP1, we constructed models of sumoylated RanGAP1 bound to both SUMO1 and Ubc9.

### Sumoylation relieves RanGAP1 autoinhibition and promotes NPC targeting

To quantitatively assess which model represents the most stable, thus most populated autoinhibited state of RanGAP1, we calculated the binding free energy between the GAP and C-terminal domains (**Fig. 5A**). M3 seems the most stable autoinhibited conformation, however, it sterically blocks access of the SUMO1/Ubc9 conjugation complex, representing sumoylation-incompetent conformational states. Although both M4 and M5 are structurally compatible with the SUMO1/Ubc9 complex, the calculated binding free energies suggest a relatively weak autoinhibitory interface in M4, but a more stable and likely more populated autoinhibited state in M5. Therefore, we selected M5 for subsequent modeling of the sumoylated RanGAP1 (denoted as a SUMO-conjugated RanGAP1/SUMO1/Ubc9 complex with an isopeptide bond between the ε-amino group of Lys524 and the C-terminal Gly97 of SUMO1). For comparison, we used the same representative conformation of M5 (**Fig. 3D**) to model the SUMO-conjugated RanGAP1/SUMO1/Ubc9 complex and conducted three independent replicate simulations.

**Fig. 5.**
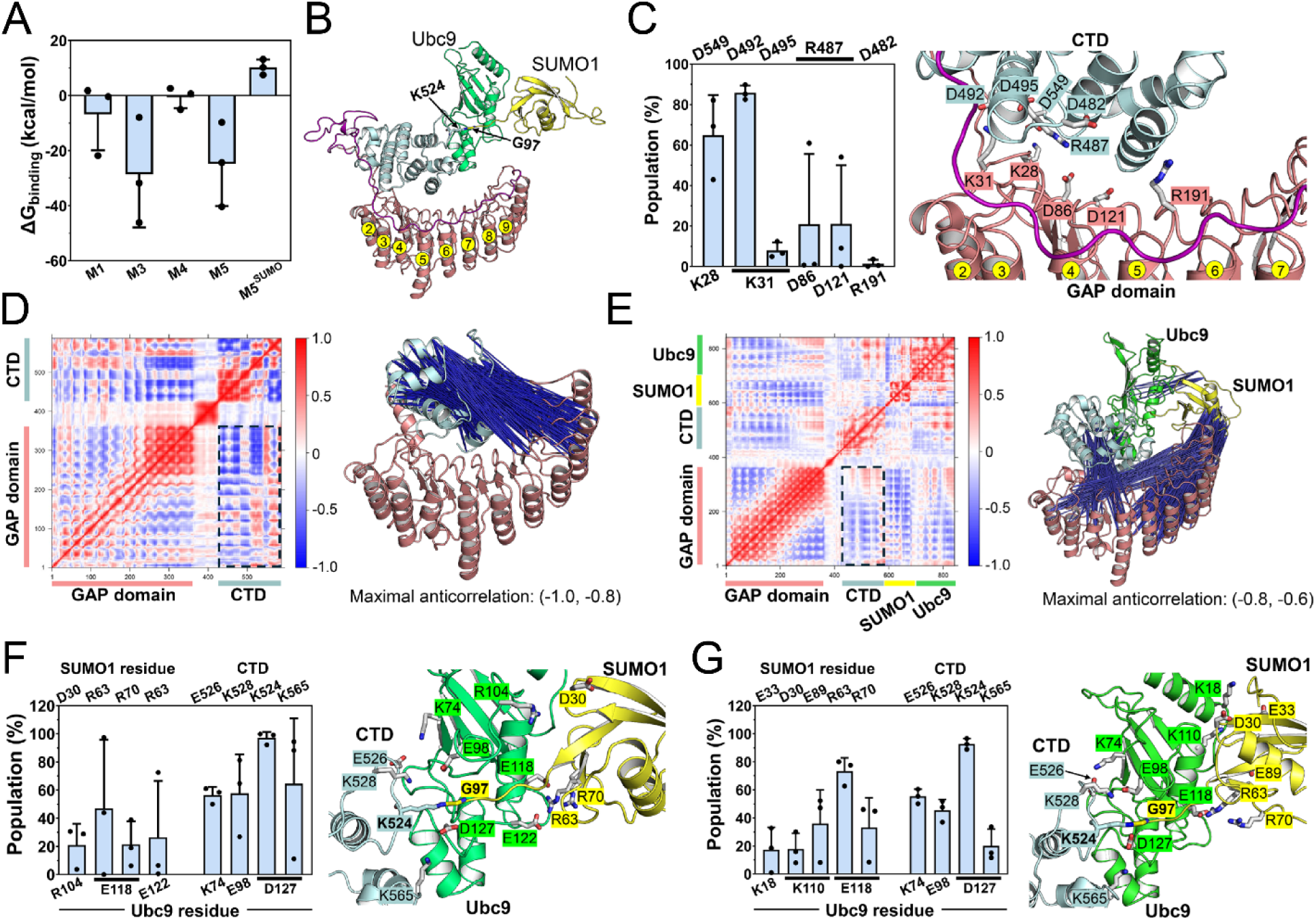
Sumoylation could allosterically relieve RanGAP1 autoinhibition. (**A**) Binding free energies between the GAP and C-terminal domains of RanGAP1 for different models. For each system, values are averaged over three independent replicate simulations. M5 exhibits the most stable autoinhibitory interaction and may represent the more populated autoinhibited conformation. In contrast, sumoylated M5 displays an unfavorable binding free energy, suggesting that sumoylation relieves RanGAP1 autoinhibition. (**B**) Representative conformation of the SUMO-conjugated RanGAP1/SUMO1/Ubc9 complex. The isopeptide bond formed between the ε-amino group of Lys524 and the C-terminal Gly97 of SUMO1 is indicated. (**C**) Complementary electrostatic interactions and their contributions to the autoinhibitory interface in the complex. (**D**) Dynamic cross-correlation matrix (DCCM) analysis indicates that there are negatively correlated motions between the GAP domain and the C-terminal domain outside of the binding interface in the autoinhibited M5. (**E**) The DCCM map also indicates that the negatively correlated motions facilitate the release of autoinhibition in the complex. (**F**) Complementary electrostatic interactions at the interface between Ubc9 and the C-terminal domain of full-length RanGAP1. (**G**) Complementary electrostatic interactions at the interface between Ubc9 and the isolated C-terminal domain of RanGAP1. The relative positioning of Ubc9 and the GAP domain is preserved in the presence and absence of the GAP domain, indicating the GAP domain has little effect on sumoylation.

**Fig. 5B** shows a representative conformation of the SUMO-conjugated RanGAP1/SUMO1/Ubc9 complex, in which the C-terminal domain appears to be dissociated from the GAP domain. This is consistent with the unfavorable binding energy between the C-terminal and GAP domains of SUMO-conjugated RanGAP1 (**Fig. 5A**). Electrostatic interactions at the autoinhibitory interface become weak compared to the autoinhibited conformation of RanGAP1 (**Fig. 5C**). Specifically, the interactions of Arg189^GAP^–Asp482^CTD^, Arg191^GAP^–Asp482^CTD^, Asp359^GAP^–Lys528^CTD^, and Asp360^GAP^–Lys530^CTD^ are diminished (**Figs. 4D** and **5C**). The newly formed Lys28^GAP^–Asp549^CTD^ and Lys31^GAP^–Asp492^CTD^ interactions help maintain the C-terminal domain at the edge (LRR 1) of the GAP domain. Although we did not observe direct dissociation of the C-terminal domain from the GAP domain throughout the simulations, the weak interaction is unlikely to maintain the autoinhibited conformation of SUMO-conjugated RanGAP1. This is further demonstrated by the results of dynamic cross-correlation matrix (DCCM) analyses for RanGAP1 and SUMO-conjugated RanGAP1. We observed both positive and negative correlations between the C-terminal and GAP domains of RanGAP1, with the strongest anticorrelated motions occurring outside the autoinhibitory interface (**Fig. 5D**). In contrast, the motions of the C-terminal and GAP domains in SUMO-conjugated RanGAP1 are predominantly anticorrelated, indicating that these two domains move in opposite directions over time (**Fig. 5E**).

To check whether the release of autoinhibition is driven by interactions between SUMO1/Ubc9 and the GAP domain, we calculated the interaction energy between SUMO1/Ubc9 and the GAP domain of SUMO-conjugated RanGAP1. For this calculation, we considered only the short-range electrostatic and van der Waals (vdW) interactions (**Fig. S2**). We observed no direct interactions between Ubc9 and the GAP domain across three replicate simulations. While SUMO1 interacted with the GAP domain, its absence in the remaining two suggests that the release of RanGAP1 autoinhibition arises from an allosteric effect rather than direct contact. Because the crystal structures of the SUMO-conjugated RanGAP1/SUMO1/Ubc9 complex (**Fig. 4E**) included only the C-terminal domain, we sought to determine how the GAP domain influences Ubc9 binding. Using the structure (PDB ID: 1Z5S) as a template, we modeled a complex containing only the C-terminal domain and performed three replicate simulations for comparison. The binding interface between Ubc9 and the C-termina domain of RanGAP1 remains largely unchanged regardless of the GAP domain’s presence (**Figs. 5F** and **5G**). Key electrostatic interactions at the interface, including Asp127^Ubc9^–Lys524^CTD^, Lys74^Ubc9^–Glu526^CTD^, and Glu98^Ubc9^–Lys528^CTD^, are preserved, facilitating SUMO1 conjugation. In contrast, at the SUMO1/Ubc9 interface, only the key interaction between Glu118^Ubc9^ and Arg63^SUMO1^ is consistently maintained. Other interactions at this interface appear unstable and distinct in the presence and the absence of the GAP domain, indicating that the GAP domain can influence SUMO1 conformation during its dissociation from the C-terminal domain of RanGAP1. This conformational flexibility may enable SUMO1 to adopt an optimal orientation for binding to the E3 ligase domain of RanBP2, thereby completing the conjugation process^43^. Taken together, these results suggest that sumoylation triggers an allosteric shift that disrupts RanGAP1 autoinhibition, allowing the SUMO-conjugated RanGAP1/SUMO1/Ubc9 complex to anchor to RanBP2.

### RanGAP1 promotes GTP hydrolysis in Ran at the cytoplasmic NPC with an arginine finger

Having characterized the sumoylation-dependent release of autoinhibition at the C-terminal domain, we next examined how the N-terminal GAP domain of RanGAP1 performs its catalytic function on the cytoplasmic side of the NPC. The ran binding domains (RBDs) of RanBP2 selectively capture Ran-GTP and position it for efficient GTP hydrolysis by the GAP domain of RanGAP1. We thus modeled the complex comprising the GAP domain of RanGAP1 (RanGAP1^GAP^), Ran-GTP, and the fourth RBD of RanBP2 (RanBP2^RBD4^) based on the available cryo-EM structure (PDB ID: 9B62) and performed three replicate simulations of the RanGAP1^GAP^/Ran-GTP/RanBP2^RBD4^ complex (**Fig. 6A**). The initial complex represents a ground OFF state, with no RanGAP1^GAP^ arginine residues contacting GTP. The binding interface between RanGAP1^GAP^ and Ran-GTP is stabilized by multiple persistent electrostatic interactions, including Arg95^Ran^–Asp273^GAP^, Lys130^Ran^–Asp245^GAP^, Lys130^Ran^–Asp273^GAP^, and Lys132^Ran^–Glu304^GAP^ (**Fig. 6B**). Notably, in one simulation, Arg191 inserts into the GTP-binding pocket and directly interacts with the γ-phosphate (**Fig. 6C** and **Fig. S3**). The difference in the Arg191–GTP distance among the three simulations suggest distinct states of RanGAP1-mediated GTP hydrolysis (**Fig. 6D**). The largest mean distance of 11 Å indicates the ground OFF state in which Arg191 is prevented from interacting with the γ-phosphate by interactions involving Tyr39 and the highly conserved Gln69 with GTP. In contrast, a distance below 3 Å corresponds to the ground ON state in which Arg191 interacts with the γ-phosphate, as observed in the Rac2/p50-RhoGAP complex^44^. Distances varying from 4 to 14 Å in one simulation indicate intermediate conformational states.

**Fig. 6.**
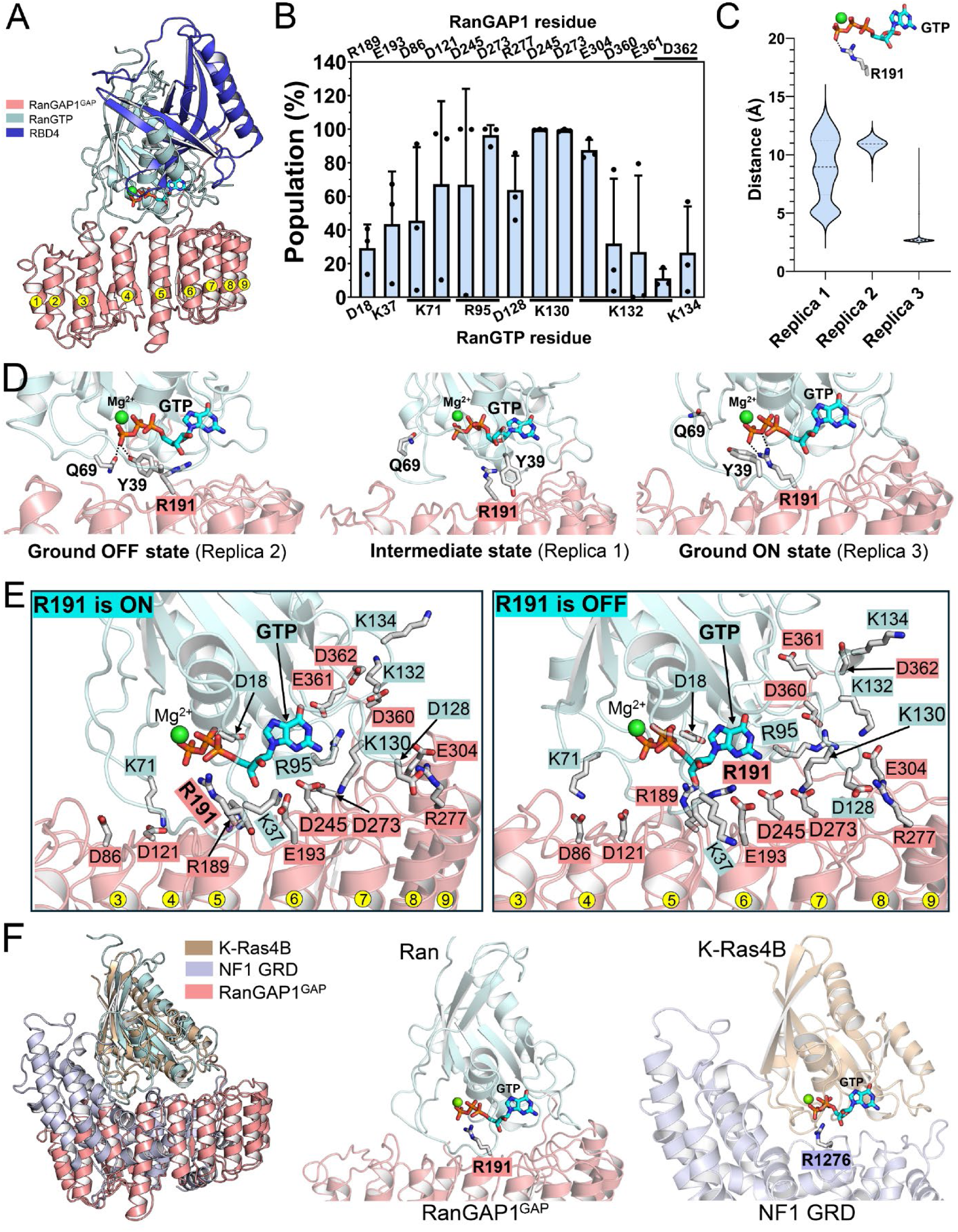
RanGAP1 mediates GTP hydrolysis in Ran at the cytoplasmic NPC with an arginine finger. (**A**) Structure of the RanGAP1^GAP^/Ran-GTP/RanBP2^RBD4^ complex, comprising the GAP domain of RanGAP1, Ran-GTP, and the fourth Ran-binding of RanBP2. The complex was constructed based on the cryo-EM structure (PDB ID: 9B62), in which Arg191 of RanGAP1 does not interact with GTP. (**B**) Complementary electrostatic interactions and their contributions to the binding between Ran-GTP and RanGAP1. The population of each pair interaction is averaged over three independent replicate simulations. (**C**) Distribution of the distance between Arg191 of RanGAP1 and GTP in three replicate simulations. Mean values are indicated by bold dashed lines. (**D**) Representative conformational states corresponding to the different distances in (C). The ground OFF-state is characterized by a large distance between Arg191 and GTP; the ground ON-state corresponds to insertion of Arg191 into the GTP-binding pocket in which it interacts with the γ-phosphate of GTP; and the intermediate state exhibits significant fluctuations in the Arg191–GTP distance. For comparison, residues Tyr39 and Gln69 are also labeled. (**E**) Interaction networks at the binding interface between Ran-GTP and the GAP domain of RanGAP1 in both ground ON and OFF states. A conserved interaction network is maintained in both states, suggesting a low energy barrier between these two states. (**F**) Structural superposition of the RanGAP1^GAP^/Ran-GTP complex with the NF1^GRD^/K-Ras4B complex. Arg1276 in the GAP-related domain (GRD) of NF1 functions as a canonical arginine finger that promotes GTP hydrolysis in K-Ras4B. A similar orientation of Arg191 in RanGAP1 relative to Ran-GTP is observed. These analyses suggest that Arg191 may act as an arginine finger.

The overall interaction network at the binding interface remains largely conserved between the ground ON and OFF states (**Fig. 6E**), suggesting that the different Arg191–GTP interaction may arise from a local conformational change, and these two states are separated by a relatively low energy barrier. To verify that the conformational transition between the OFF and ON states is energetically accessible, we performed two independent accelerated MD (aMD) simulations of the RanGAP1^GAP^/Ran-GTP/RanBP2^RBD4^ complex. As in our prior study of M-Ras^45^, aMD with dihedral boosting enhances conformational sampling by lowering torsional energy barriers, facilitating transitions involving side chain and local structural rearrangements. Arg191 rapidly inserts into the GTP-binding pocket and forms stable interactions with the γ-phosphate. These observations suggest that the transition is energetically accessible and involves a low barrier (**Fig. S4**).

To determine whether Arg191 functions as an arginine finger and follows the canonical mechanism of GAP-mediated GTP hydrolysis observed in Ras and Rho GTPases, we compared the ground ON state conformations of RanGAP1^GAP^/Ran-GTP with those of the GAP-related domain of neurofibromin (NF1^GRD^) in complex with K-Ras4B from our previous study^39^. We superimposed the representative conformation of RanGAP1^GAP^/Ran-GTP with that of NF1^GRD^/K-Ras4B and observed a similar orientation of Arg191 in RanGAP1^GAP^ and the arginine finger Arg1276 in NF1^GRD^ relative to GTP (**Fig. 6F**). In K-Ras4B, Gln61 is an important catalytic residue and contributes to GTP hydrolysis by orienting the nucleophilic water molecule toward the γ-phosphate of GTP. In the ground ON state of NF1^GRD^/K-Ras4B complex, the average distance between Gln61 and the γ-phosphate of GTP is 7.4 Å, and a comparable distance of 7.8 Å was observed between the equivalent Gln69 and the γ-phosphate of GTP in the RanGAP1^GAP^/Ran-GTP complex (**Fig. S5**). Also, in our previous study, we identified the Glu63–Arg1391 salt bridge at the binding interface that stabilizes the conformation of Gln61 in K-Ras4B. Similarly, Lys71 of Ran forms salt bridges with Asp86 or Asp121 of RanGAP1^GAP^ at the binding interface, modulating the orientation of Gln69^Ran^ (**Figs. 6B**, **6E**, and **S5**). In the ground ON state, Lys71^Ran^ preferentially interacts with Asp121^GAP^ that is closer to GTP, facilitating proper positioning Gln69^Ran^ for catalysis. In contrast, in the ground OFF state, Lys71^Ran^ preferentially interacts with Asp86^GAP^, allowing greater flexibility of Gln69 with respect to GTP (**Figs. 6E** and **S5**). Collectively, these results suggest that Arg191 satisfies the structural requirements for a canonical arginine finger and clarify why Gln69^Ran^ is highly conserved.

### RanGAP1 promotes GTP hydrolysis in Ran in the cytosol as a backup mechanism

Although RanGAP1 preferentially localizes to the cytoplasmic side of the NPC through sumoylation-dependent interaction with RanBP2, it is also present in the cytosol. Similarly, while Ran-GTP is predominantly localized in the nucleus, a fraction of Ran-GTP diffuses into the cytosol, where Ran-GTP forms a ternary complex with RanBP1 and RanGAP1 to promote GTP hydrolysis. Mechanistic understanding of the RanGAP1^GAP^/Ran-GTP/RanBP1 complex is largely inferred from studies of Ran-GTP bound to RanBP1 (human) and Rna1p (yeast)^30^. Although the LRR fold of the GAP domain is highly conserved between human and yeast, their sequence identity is low (∼30–36%) (**Fig. S6**). Moreover, Rna1p lacks the C-terminal domain present in RanGAP1 and therefore does not require sumoylation for its localization and function. The different structural contexts between human and yeast suggest that RanGAP1 may function by a different molecular mechanism. To investigate how RanGAP1 promotes GTP hydrolysis in the cytosol, we modeled the RanGAP1^GAP^/Ran-GTP/RanBP1 complex based on two available experimental structures (PDB IDs: 1K5D and 9B62, see Methods for details) (**Fig. 7A**), and performed three replicate simulations.

**Fig. 7.**
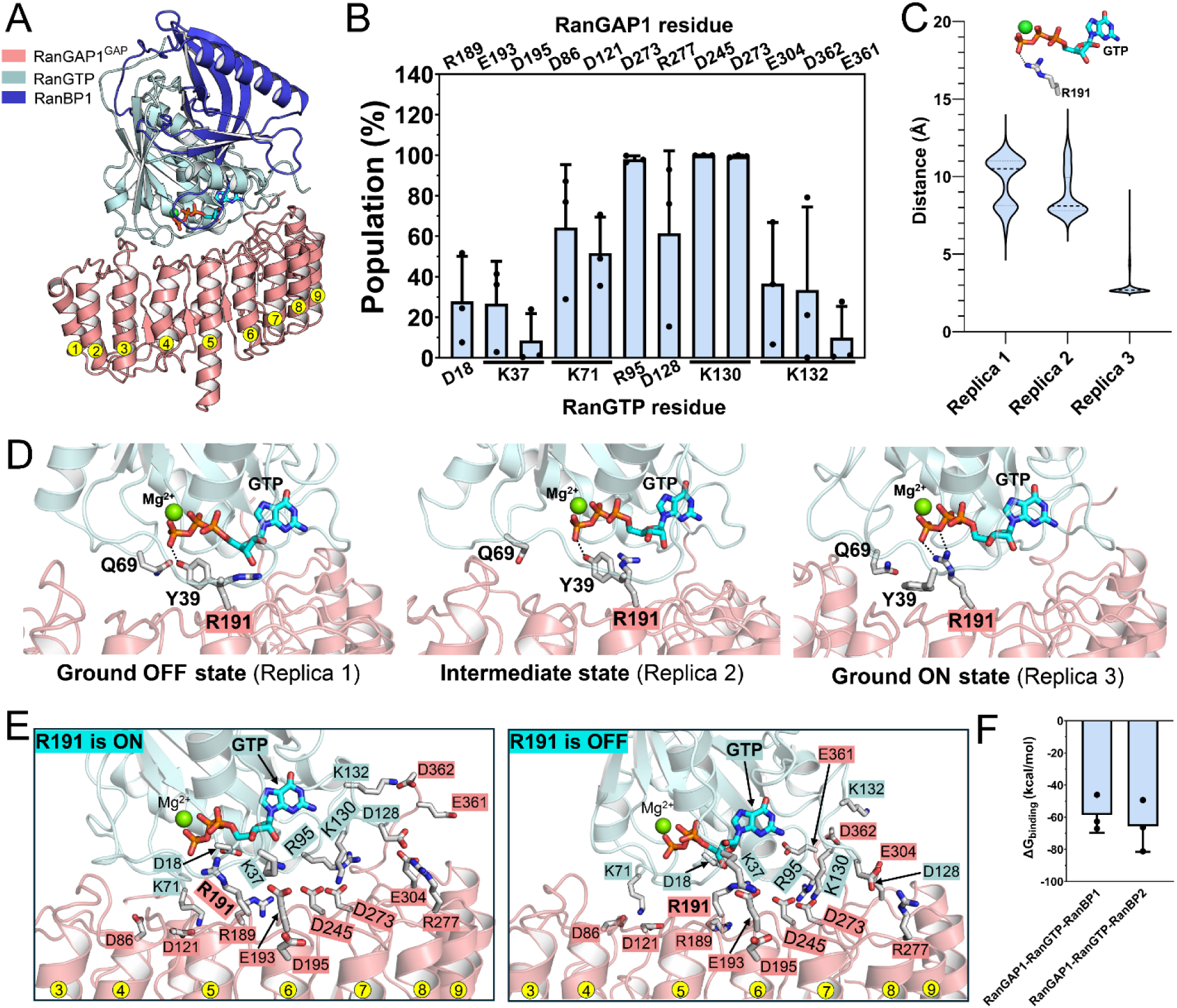
RanGAP1 accelerates GTP hydrolysis in Ran in the cytosol. (**A**) Structure of the RanGAP1^GAP^/Ran-GTP/RanBP1 complex, comprising the GAP domain of RanGAP1, Ran-GTP, and RanBP1. In this complex, Arg191 of RanGAP1 does not interact with GTP initially. (**B**) Complementary electrostatic interactions and their contributions to the binding between Ran-GTP and RanGAP1. The population of each pair interaction is averaged over three independent replicate simulations. (**C**) Distribution of the distance between Arg191 of RanGAP1 and GTP in three replicate simulations. Mean values are indicated by bold dashed lines. (**D**) Representative conformational states corresponding to the different distances in (C). The ground OFF-state is characterized by a large distance between Arg191 and GTP; the ground ON-state corresponds to insertion of Arg191 into the GTP-binding pocket in which it interacts with the γ-phosphate of GTP; and the intermediate state exhibits large fluctuations in the Arg191–GTP distance. For comparison, residues Tyr39 and Gln69 are also labeled (**E**) Interaction networks at the binding interface between Ran-GTP and the GAP domain of RanGAP1 in both ground ON and OFF states. A conserved interaction network is maintained in both states, suggesting a low energy barrier between these two states. (**F**) Binding free energies for the RanGAP1^GAP^/Ran-GTP/RanBP1 and RanGAP1^GAP^/Ran-GTP/RanBP2^RBD4^ complexes. Values are averaged over three independent replicate simulations. RanGAP1-mediated GTP hydrolysis in Ran in the cytosol serves as a backup mechanism for maintaining the Ran-GTP gradient across the NPC.

We first characterized the binding interface between Ran-GTP and RanGAP1^GAP^ (**Fig. 7B**) and observed electrostatic interactions similar to those demonstrated in the RanGAP1^GAP^/Ran-GTP/RanBP2^RBD4^ complex (**Fig. 6B**). In the complex, Arg95^Ran^ displays a more specific interaction with Asp273^GAP^, whereas Arg95^Ran^ interacts with Asp273^GAP^ and Asp245^GAP^ as well. In one of the three replicate simulations, we also observed that Arg191 inserts into the GTP-binding pocket and interacts with the γ-phosphate (**Figs. 7C** and **S7**). Correspondingly, the three replicate simulations represent the ground OFF, intermediate, and ground ON states based on the Arg191–GTP distance (**Fig. 7D**). The conserved interaction network between the ground ON and OFF states suggests that they are separated by a low energy barrier (**Fig. 7E)**. The distances between Gln69^Ran^ and the γ-phosphate of GTP are all below 8 Å (**Fig. S7**), and the neighboring Lys71^Ran^ interacts with Asp86^GAP^ and Asp121^GAP^ (**Fig. 7B**), facilitating optimal positioning of Gln69^Ran^ for catalysis. These structural features support the role of Arg191^GAP^ as an arginine finger in the RanGAP1^GAP^/Ran-GTP/RanBP1 complex. Comparable binding free energies between Ran-GTP and RanGAP1^GAP^ are obtained for both RanGAP1^GAP^/Ran-GTP/RanBP1 and RanGAP1^GAP^/Ran-GTP/RanBP2^RBD4^ complexes (**Fig. 7F**). These results are consistent with previous experimental studies that demonstrated a common role of RanBP1 and RanBP2 as Ran-GTP-binding cofactors in facilitating RanGAP1-mediated GTP hydrolysis^23,31,46–49^, and further reveal that Arg191^GAP^ functions as a canonical arginine finger in this process. The RanGAP1^GAP^/Ran-GTP/RanBP1 complex may serve as a backup pathway for GTP hydrolysis in the cytosol, complementing the NPC-associated RanGAP1^GAP^/Ran-GTP/RanBP2 complex. In addition to the more populated autoinhibited conformation, RanGAP1 may also adopt less populated open conformation in which there are no interactions between C-terminal and GAP domains (**Fig. 3E**). Binding of Ran-GTP/RanBP1 with such open RanGAP1 could lead to stable RanGAP1^GAP^/Ran-GTP/RanBP1 complex and promote GTP hydrolysis in Ran, independent of sumoylation (**Fig. 7F**). After RanGAP-catalyzed hydrolysis of GTP to GDP, the C-terminal region of Ran adopts a closed conformation by associating with its G-domain, preventing the interaction with Ran-binding proteins^50^.

## Discussion

Nucleocytoplasmic transport is highly efficient^51–53^, with overlapping nuclear import and export paths^54^. Single-molecule fluorescence microscopy measurement of a model protein substrate indicated that NPCs could transport at least 10 substrate molecules simultaneously, and at least 1,000 molecules per NPC per second^55^. This requires rapid and tightly coordinated assembly and disassembly of transport complexes within crowded cellular environments. At the cytoplasmic face of the NPC, multiple factors, including Ran-GTP, transport receptors like CRM1, Ran-binding proteins, and RanGAP1, need to act efficiently to ensure directional transport^56,57^. In this context, the structural organization and regulation of RanGAP1 is particularly important. It consists of an N-terminal GAP domain and a C-terminal sumoylation domain separated by a disordered linker (∼70 residues), raising the question of how these spatially distant functional modules are coordinated to achieve efficient catalysis.

We propose a model in which RanGAP1 adopts an autoinhibited conformation that is transiently stabilized by interactions between the GAP and C-terminal domains (**Fig. 4**). In the crowded cellular environment, autoinhibition of RanGAP1 helps maintain structural compactness and coordinates its dual functional roles within the SUMO-conjugated RanGAP1/SUMO1/Ubc9/RanBP2 complex. In the autoinhibited state, the GAP and C-terminal domains are spatially confined, facilitating coupling between sumoylation-dependent E3 ligase activity and subsequent GTP hydrolysis in Ran. Upon sumoylation, RanGAP1 is recruited to RanBP2 at the cytoplasmic filaments of the NPC, forming the RanGAP1/SUMO1/Ubc9/RanBP2 complex that catalyzes SUMO1 conjugation and acts as a disassembly machine for CRM1-dependent export complexes^28^. Consistent with this mechanism, our results suggest that sumoylation could allosterically relieve autoinhibition (**Fig. 5**), enabling the SUMO-conjugated RanGAP1/SUMO1/Ubc9 complex to associate with RanBP2 and allowing the GAP domain to efficiently hydrolyze GTP in Ran, driving export complex disassembly. Autoinhibitory regulation is also observed in other components of the nucleocytoplasmic transport machinery. For example, importin-α adopts an autoinhibited conformation in which its N-terminal importin-β–binding domain occupies the NLS-binding site to prevent cargo binding in the nucleus^58^. This inhibition is relieved upon binding to importin-β in the cytoplasm, allowing the importin-α/importin-β heterodimer to bind and transport a cargo through the NPC into the nucleus^59^. Together, these results indicate that autoinhibition may represent a common strategy in nucleocytoplasmic transport.

Our analyses of the RanGAP1^GAP^/Ran-GTP/RanBP2^RBD4^ and RanGAP1^GAP^/Ran-GTP/RanBP1 complexes provide a structural basis for the distinct mechanisms of release of RanGAP1 autoinhibition and GTP hydrolysis in Ran in different contexts (**Figs. 6** and **7**). The RanBP2-associated RanGAP1 is the primary platform for GTP hydrolysis. Binding of a RanBP2 RBD with the CRM1/Ran-GTP/cargo complex induces cargo release and formation of a CRM1/Ran-GTP/RanBP2^RBD^ intermediate, which then dissociates to produce a Ran-GTP/RanBP2^RBD^ subcomplex. RanBP2 contains four RBDs, which increases the local concentration of Ran-GTP at the cytoplasmic filaments of the NPC. The proximity between the sumoylated C-terminal domain and the GAP domain of RanGAP1 on RanBP2 provides a spatial advantage and allows the GAP domain to access the Ran-GTP/’RanBP2^RBD^ subcomplex and initiate hydrolysis efficiently.

During nuclear export, the CRM1/Ran-GTP/cargo complex translocates through the NPC to the cytoplasmic side where RanBP2 and RanBP1 promote RanGAP1-catalyzed hydrolysis, leading to complex disassembly and cargo release. This process results in scarcity of CRM1/Ran-GTP in the cytosol. Similarly, during nuclear import, the importin-α**/**importin-β**/**cargo complex is disassembled in the nucleus upon Ran-GTP binding to importin-β, leading to cargo release. Importin-β exits the nucleus in complex with Ran-GTP^60^. Thus, both export and import pathways generate Ran-GTP-bound receptor complexes^61^. To maintain the Ran-GTP gradient across the NPC, characterized by a low concentration of Ran-GTP in the cytosol, cytosolic RanBP1 binds to CRM1/Ran-GTP and importin-β/Ran-GTP complexes, promoting release of the Ran-GTP/RanBP1 intermediate that is subsequently hydrolyzed by RanGAP1^62,63^. Our analyses of RanGAP1^GAP^/Ran-GTP/RanBP1 complex suggest that Ran-GTP/RanBP1 could bind to the less populated open conformation of RanGAP1 in a sumoylation-independent manner (**Fig. 7**), and follows a similar GTP hydrolysis mechanism observed in the RanGAP1^GAP^/Ran-GTP/RanBP2^RBD4^ complex (**Fig. 6**). Thus, the RanBP1-mediated pathway acts as a backup mechanism, ensuring efficient GTP hydrolysis in Ran, not captured by RanBP2 at NPC. **Fig. 8** summarizes these two distinct pathways.

**Fig. 8.**
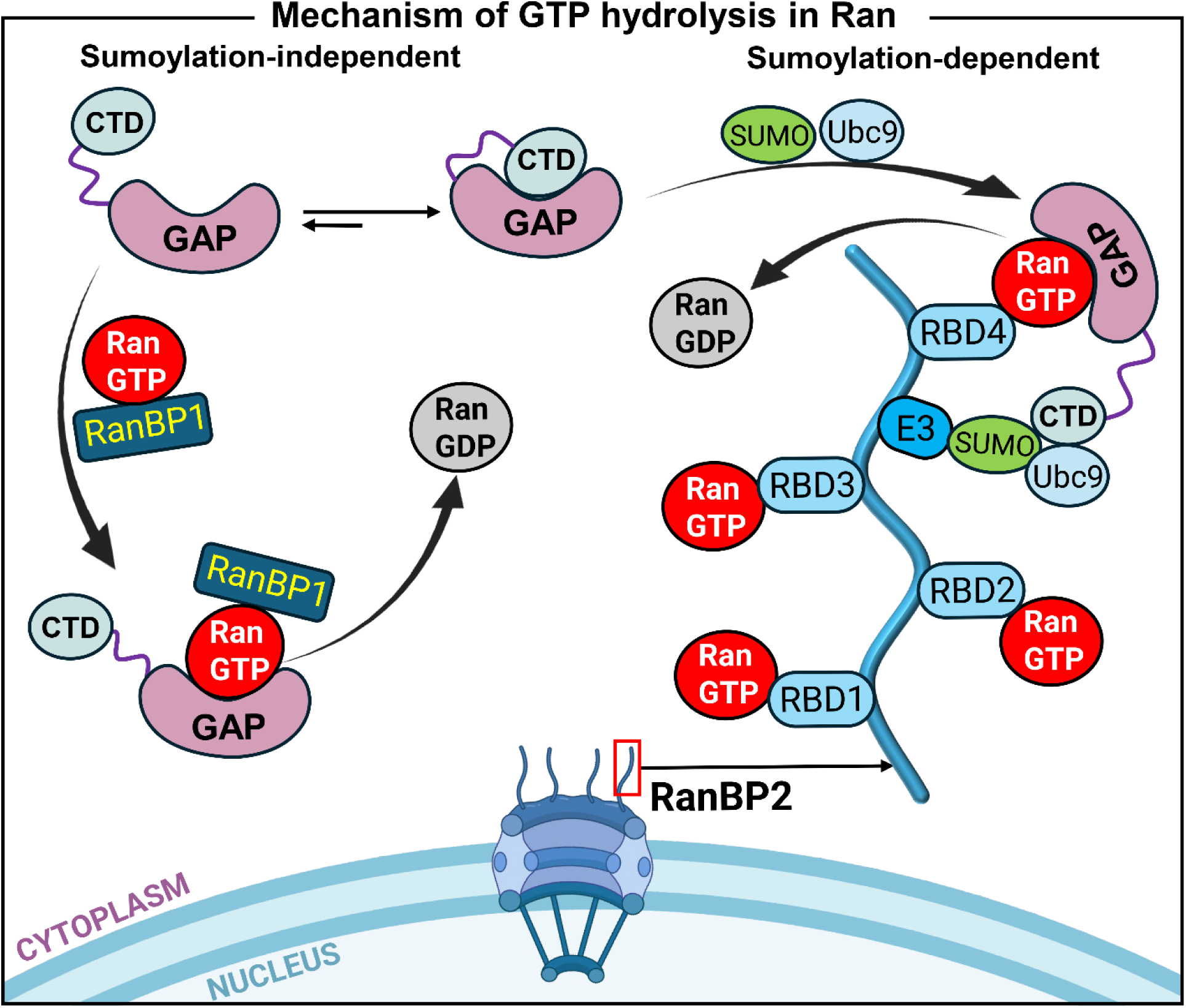
Schematic summary of two distinct mechanisms for the relief of RanGAP1 autoinhibition and subsequent GTP hypothesis at the cytoplasmic filaments of RanBP2 (primary pathway) or in the cytoplasm (backup pathway). RanGAP1 exists in an equilibrium between a predominantly populated autoinhibited conformation and a less populated open conformation. In the primary pathway, the export complex is disassembled upon interaction with the Ran-binding domains (RBDs) of RanBP2, leading to cargo release. The binding of Ran-GTP to multiple RBDs contributes to its local enrichment at RanBP2. RanGAP1 relieves its autoinhibition upon sumoylation of its C-terminal domain and subsequent association with RanBP2. The resulting SUMO-conjugated RanGAP1/SUMO1/Ubc9/RabBP2 complex functions as a SUMO E3 ligase that facilitates completion of the sumoylation process. Concurrently, the GAP domain of RanGAP1 engages Ran-GTP bound to the RBDs (illustrated with RBD4) and promotes GTP hydrolysis. In the cytoplasm, RanBP1 further stimulates RanGAP1-catalyzed GTP hydrolysis and contributes to disassembly of the export complex, resulting in Ran-GTP/RanBP1 complexes. In the backup pathway, the Ran-GTP/RanBP1 complex interacts with the GAP domain of RanGAP1 in less populated open conformation and enable GTP hydrolysis in the cytosol. This process maintains low cytosolic Ran-GTP levels. In both pathways, RanGAP1-mediated hydrolysis involves Arg191 that may act as an arginine finger. Part of this graphic is created with BioRender.

In both RanBP1- and RanBP2-facilitated RanGAP1-mediated GTP hydrolysis, our simulations show that Arg191 of RanGAP1 inserts into the GTP-binding pocket and interacts with the γ-phosphate, acting as a canonical arginine finger, similar to that in the NF1^GRD^/K-Ras4B complex (**Figs. 6** and **7**). This finding differs from previous studies that used Rna1p and found no arginine residue involved in GTP hydrolysis. For instance, earlier mutational studies showed that conserved arginine residues Arg182 and Arg184 in budding yeast (*Saccharomyces cerevisiae*) Rna1p (equivalent to Arg189 and Arg191 in human RanGAP1) had no effect on Rna1p activity^64^. Later, the crystal structures of the complex formed by human Ran (GppNHp-bound), human RanBP1, and fission yeast (*Schizosaccharomyces pombe*) Rna1p in the ground state and a transition-state analogue indicated that GTP hydrolysis does not require an arginine finger^30^.

The yeast Rna1p shares low sequence identity with the GAP domain of human RanGAP1 (**Fig. S6**) and is predominantly cytosolic due to the absence of the sumoylation domain. These sequence and localization differences raise the question of whether the same catalytic mechanism applies to the human system. The recent cryo-EM structure captures a ground OFF state conformation of the RanGAP1/Ran^Q69L^-GTP/RanBP2 complex, in which Arg191 does not insert into the GTP-binding pocket. However, consistent with our previous simulations of the NF1^GRD^/K-Ras4B complex, in which the arginine finger Arg1276 interacts with GTP from an initial OFF state conformation, our current simulations show that Arg191 can also interact with GTP from the starting ground OFF state conformation in both RanGAP1^GAP^/Ran-GTP/RanBP1 and RanGAP1^GAP^/Ran-GTP/RanBP2^RBD4^ complexes. Note that the NF1 R1276A mutant reduces its GAP activity by about 1000-fold^65^. Thus, the presence of Arg191 in human RanGAP1 may provide an evolutionary advantage that enhances the GTP hydrolysis efficiency in human nuclear transport, given the relatively low intrinsic hydrolysis rate of Ran compared with other GTPases^66^. The mechanistic divergence between conserved protein fold and distinct mechanisms has also been observed in RanBP1. Although structurally conserved between fungi and animals, RanBP1 employs different mechanisms for cargo dissociation and nuclear export^67^. Thus, conserved protein folds do not necessarily imply conserved molecular mechanisms. Together, our results highlight a catalytic role of Arg191 in RanGAP1-mediated GTP hydrolysis and provide a testable hypothesis that can be experimentally validated through site-directed mutagenesis and functional assays.

In conclusion, we establish how human RanGAP1 regulates sumoylation and GTP hydrolysis in Ran at the cytoplasmic filament of NPC during nucleocytoplasmic transport. Our results suggest that RanGAP1 adopts a dynamic, autoinhibited conformation, and this autoinhibition can be allosterically relieved upon sumoylation and recruitment to RanBP2, allowing RanGAP1 to interact with Ran-GTP and facilitate disassembly of export complexes. The identification of Arg191 as an arginine finger suggests a divergence in catalytic mechanism between human and yeast. These findings underscore the importance of conformational regulation and sumoylation modification in controlling complex cellular processes in a crowded cellular environment.

## Methods

### Structural modeling of full-length RanGAP1 in autoinhibited conformations

The full-length human RanGAP1 (UniProt identifier: P46060) is a 587-residue protein, comprising an N-terminal GAP domain, a disordered linker, and a C-terminal domain (**Fig. 1B**). No intradomain interactions are predicted by AlphaFold2 (**Fig. S8**)^68^. To model the autoinhibited RanGAP1 in which the C-terminal associates with the GAP domain, we extracted the two domains from the recent cryo-EM structure (PDB ID: 9B62)^38^ and performed Rosetta protein–protein docking^69^. The modeling protocol follows our previous work on autoinhibited conformations of 3-phosphoinositide-dependent kinase 1 (PDK1)^70^. We performed two independent Rosetta docking, resulting in a total of 20,000 decoys for the complex. We ranked these decoys based on the Rosetta’s total score’(T_sc) and interface score (I_sc) to select the overall stable complexes with favorable domain interactions. The top five models were selected for further refinement. We then modeled the disordered linker using the cyclic coordinate descent method and refined using the kinematic closure method within the Rosetta loop modeling module^71–73^. Among the generated linker conformations, we selected those adopting predominantly random coil structures with minimal contacts with either domain. The resulting autoinhibited conformations of RanGAP1 are shown in **Fig. 2**.

### Structural modeling of sumoylated RanGAP1 and RanGAP1/Ran-GTP/RanBP complexes

The SUMO E2 enzyme Ubc9 exhibits high affinity for the C-terminal domain of RanGAP1 during SUMO1 transfer. To satisfy the specific spatial arrangement among RanGAP1, SUMO1 and Ubc9, we modeled the SUMO-conjugated RanGAP1 by superposing the C-terminal domain of the autoinhibited RanGAP1 onto the crystal structure of RanGAP1 C-terminal domain in complex with SUMO1 and Ubc9 (PDB ID: 1Z5S)^74^. In the SUMO-conjugated RanGAP1/SUMO1/Ubc9 complex, an isopeptide bond is built between the ε-amino group of the sumoylation site Lys524 of RanGAP1 and the C-terminal Gly97 of SUMO1. The corresponding force filed parameters were taken from our previous study^75^.

To model the interaction of Ran-GTP with RanGAP1 in the presence of RanBP1 or RanBP2, we constructed two complexes using the GAP domain of RanGAP1: RanGAP1^GAP^/Ran-GTP/RanBP1 and RanGAP1^GAP^/Ran-GTP/RanBP2^RBD4^. The RanGAP1^GAP^/Ran-GTP/RanBP2^RBD4^ was modeled based on the cryo-EM structure (PDB ID: 9B62)^38^, which includes the GAP domain of RanGAP1, the full-length Ran-GTP, and the fourth RBD of RanBP2. To build the RanGAP1^GAP^/Ran-GTP/RanBP1 complex, we extracted the Ran-GTP/RanBP1 structure from the crystal structure of GppNHp-bound Ran in complex with RanBP1 and Rna1p complex (PDB ID: 1K5D)^30^, and then superimposed it onto the RanGAP1^GAP^/Ran-GTP/RanBP2^RBD4^ model to generate the human complex.

### Atomistic MD Simulations Protocols

All MD simulations were carried out using NAMD 2.14^76,77^. The simulation protocols were similar to those used in our previous works^78^. Each protein structure was represented using the updated and modified version of the CHARMM36m force field parameters^79^. Each system was solved in a cubic water box filled with TIP3P water molecules, with a minimum distance of 12 Å between protein and the edge of the water box, and neutralized with 100 mM NaCl. Temperature and pressure were maintained at 310 K and 1 atm using Langevin dynamics and the Nosé–Hoover Langevin piston method, respectively^80,81^. Long-range electrostatic interactions were treated with the particle mesh Ewald (PME) method, using a grid spacing of 1.0 Å and an interpolation order of 6^82^. The vdW interactions were calculated using a switching function with a twin cutoff of 10 and 12 Å. Bonds involving hydrogen atoms were constrained using the SHAKE algorithm, allowing an integration time step of 2 fs^83^. Each system was energy minimized for 20,000 steps and equilibrated for 5 ns in the NVT ensemble (310 K) with harmonic restraints applied to Cα atoms. Production simulations were then carried out in the NPT ensemble (1 atm, 310 K) without restraints for 1,000 ns. For each system, we performed three independent replicate simulations. Analyses were performed based on the second half of each trajectory.

### Conformational dynamic analysis

Dynamic cross-correlation matrix (DCCM) analysis was performed using the Bio3D program^84,85^ to quantify correlated motions between residues, i.e., the extent to which atomic fluctuations are correlated throughout the simulation. The dynamical cross-correlation *C_ij_* between the *i*th and *j*th Cα atoms is defined as:

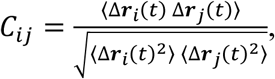

where ***r****_i_(t)* represents the position vector of the *i*th atom at time *t*, 〈·〉 denotes the time average, and Δ***r****i*(*t*) = ***r****i*(*t*) − 〈***r****i*(*t*)〉 is the displacement from the mean position. The *C_ij_* value ranges from -1 to 1. Positive values indicate correlated motion, where residues move in the same direction, whereas negative values indicate anti-correlated motion, where residues move in opposite directions. Values close to zero correspond to uncorrelated motions.

### Binding free energy calculation

Binding free energies were calculated using the molecular mechanics/generalized Born surface area (MM/GBSA) method^86,87^. The average binding free energy, ⟨Δ*G_b_*⟩, is expressed as the sum of the gas-phase molecular mechanics energy, ⟨Δ*E*_M_⟩, the solvation free energy contribution, ⟨Δ*G*_sol_⟩, and the entropy term, −*T*Δ*S*:

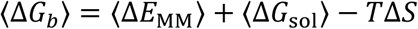

where 〈·〉 denotes averaging over MD trajectories. The gas-phase molecular mechanics energy is given by:

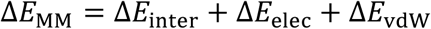

including bonded interactions, electrostatic interactions, and van der Waals contributions. The solvation free energy is decomposed into electrostatic and nonpolar components:

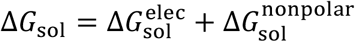

The electrostatic solvation energy was calculated using the generalized Born (GB) model^88^. The nonpolar contribution was estimated as:

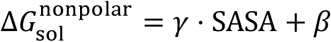

where SASA is the solvent-accessible surface area^89^, with surface tension parameter *γ* = 0.00542 kcal·mol^-1^·Å^-2^ and offset *β* = 0.92 kcal·mol^-1^.

The entropy contribution, *T*Δ*S*, was estimated using normal mode analysis^90^. Binding free energy changes upon complex formation were calculated as:

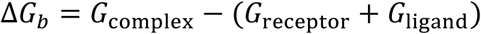

where the receptor corresponds to the GAP domain of RanGAP1, and the ligand represents either the C-terminal domain of RanGAP1 or RanGTP, depending on the specific complex analyzed. Binding free energy was computed from conformational snapshots extracted from MD trajectories. A total of 2,500 snapshots from the second half of each simulation were used to calculate the gas-phase and solvation energy terms, while 50 snapshots were used for entropy estimation due to heavy computational cost. Final Δ*G_b_* values were averaged over three independent replicate simulations. All MM/GBSA calculations were performed using the mm_pbsa.pl script implemented in Amber22 with default parameters^91^. It should be noted that absolute binding free energies obtained from MM/GBSA calculations may be much lower than the experimental data; therefore, only the relative values are reasonable^87^.

## Author contributions

H. J. and R. N. conceived the study. L. X. performed the simulations, analyzed data, and wrote the first draft of the manuscript. H. J. and R. N. commented on and revised the manuscript. All authors read and approved of the final manuscript.

## Acknowledgements

This Research was supported by the Cancer Innovation Laboratory, Center for Cancer Research, National Cancer Institute, National Institutes of Health Intramural Research Program project number ZIA BC 010441 and federal funds from the National Cancer Institute, National Institutes of Health, under contract HHSN261201500003I. The contributions of the NIH authors were made as part of their official duties as NIH federal employees, are in compliance with agency policy requirements, and are considered Works of the United States Government. However, the findings and conclusions presented in this paper are those of the authors and do not necessarily reflect the views of the NIH or the U.S. Department of Health and Human Services. The calculations had been performed using the high-performance computational facilities of the Biowulf PC/Linux cluster at the National Institutes of Health, Bethesda, MD (https://hpc.nih.gov/).

## Data availability

The research data presented in this study are documented in the paper. The representative trajectories generated from MD simulations and one trajectory movie showing interaction between Arg191 and GTP were deposited to Zenodo and can be accessed via the link: https://doi.org/10.5281/zenodo.19697122.

