## Supplemental Figure for "The structural basis of RanGAP1 regulation and catalysis in nuclear transport"

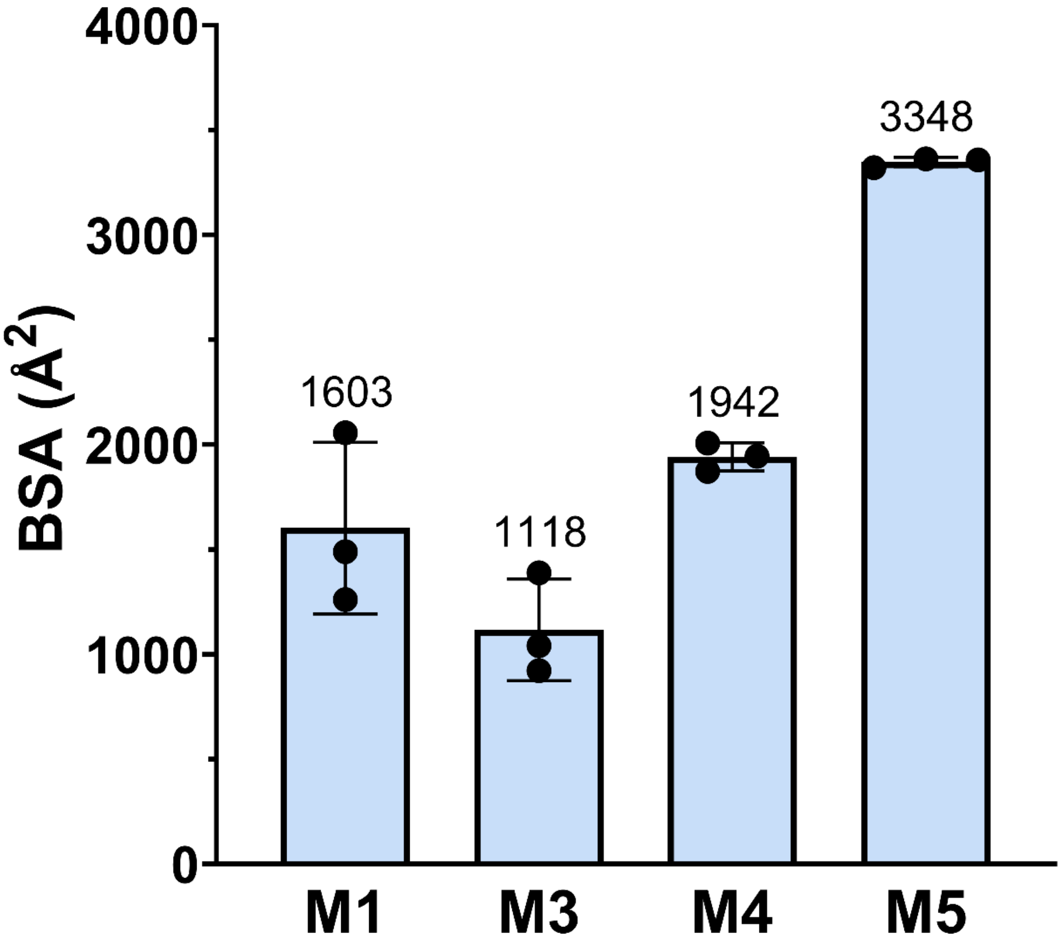


**Fig. S1. Buried surface areas of autoinhibited RanGAP1 models.** Values are averaged over three independent replicate simulations for each system. M5 exhibits the largest buried surface area, suggesting a stronger association between the C-terminal and GAP domains.

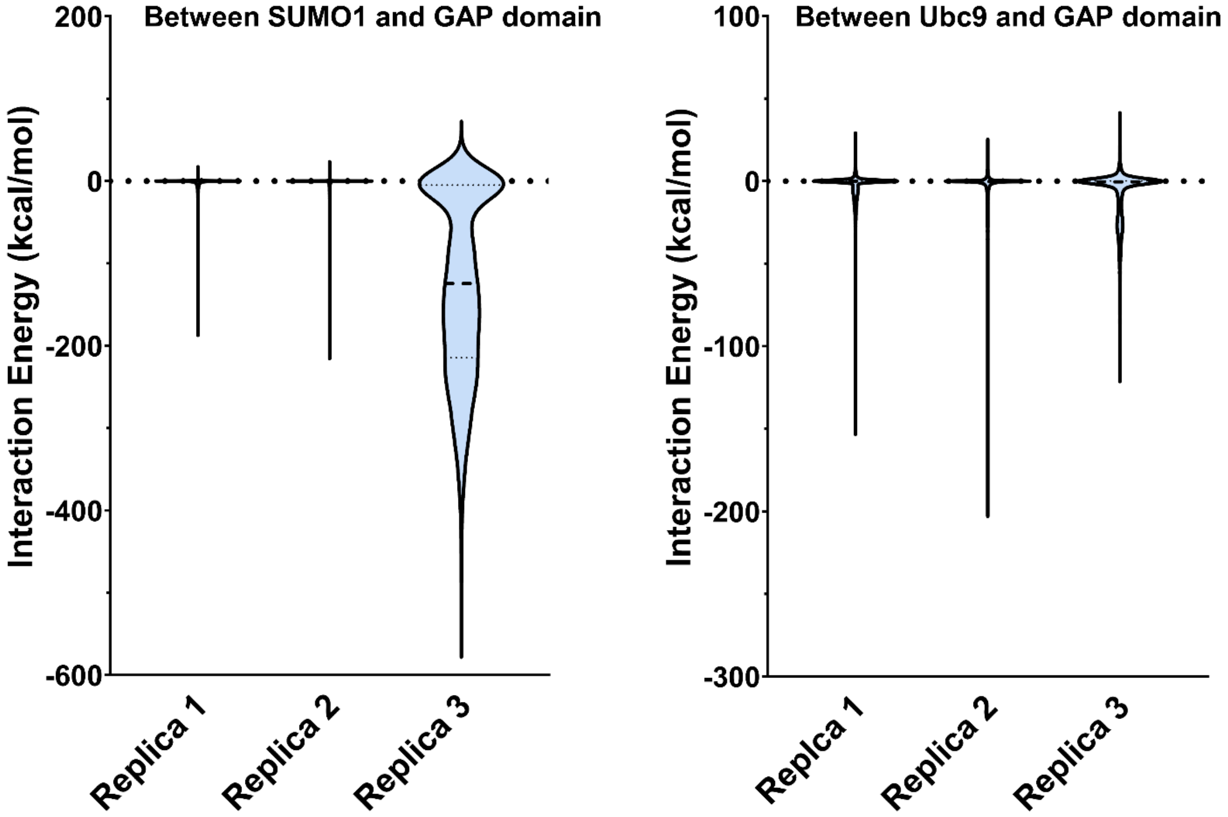


**Fig. S2. Interaction energies between SUMO1 (left) or Ubc9 (right) and the GAP domain of sumoylated RanGAP1 in the SUMO-conjugated RanGAP1/SUMO1/Ubc9 complex.** Only short-range electrostatic and van der Waals interactions are considered. Values are averaged over three independent replicate simulations for each system. No direct interactions between Ubc9 and the GAP domain were observed in any of the three simulations, whereas direct interactions between SUMO1 and the GAP domain were observed in only one replicate simulation. These results suggest that the release of RanGAP1 autoinhibition in the remaining two simulations is not driven by direct interactions between Ubc9/SUMO1 and the GAP domain, but instead arises from an allosteric effect.

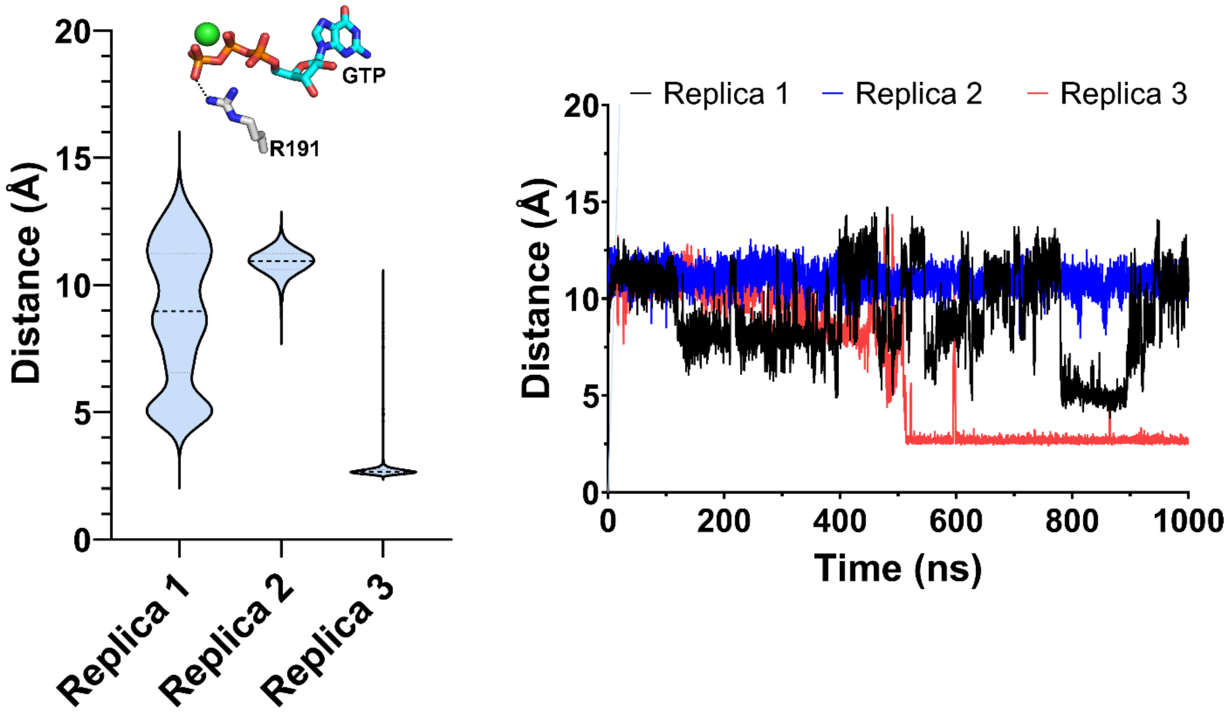


**Fig. S3. Distance between Arg191 and GTP in the RanGAP1$^{GAP}$/Ran-GTP/RanBP2$^{RBD4}$ complex.** (*Left*) Violin plots obtained from the second halves of three independent replicate simulations. (*Right*) Time evolution of the distance between Arg191 and the γ-phosphate of GTP in each simulation.

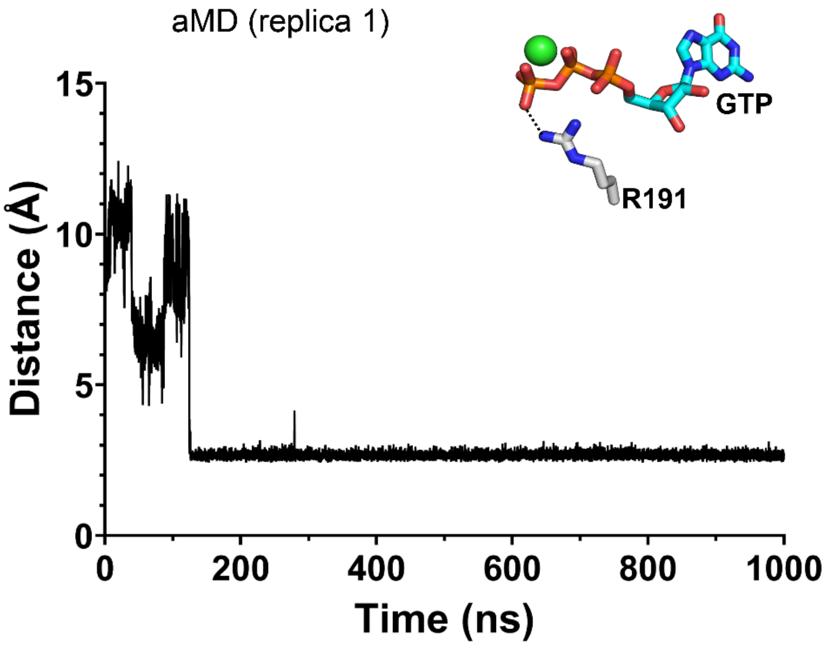


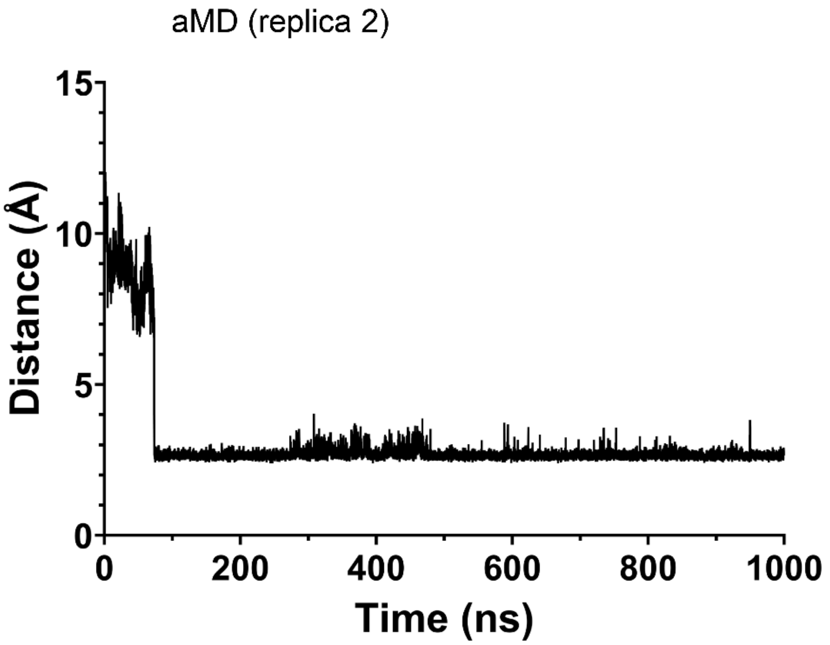


**Fig. S4. Time evolution of the distance between Arg191 and the γ-phosphate of GTP in two accelerated MD (aMD) simulations**. In the aMD simulation, a dihedral boosting was applied to enhance conformational sampling by lowering torsional energy barriers. Compared with replica 3 in Fig. S3, Arg191 rapidly inserts into the GTP-binding pocket and forms stable interactions with the γ-phosphate during the simulation. This observation confirms that the conformational transition is energetically accessible and involves a low barrier.

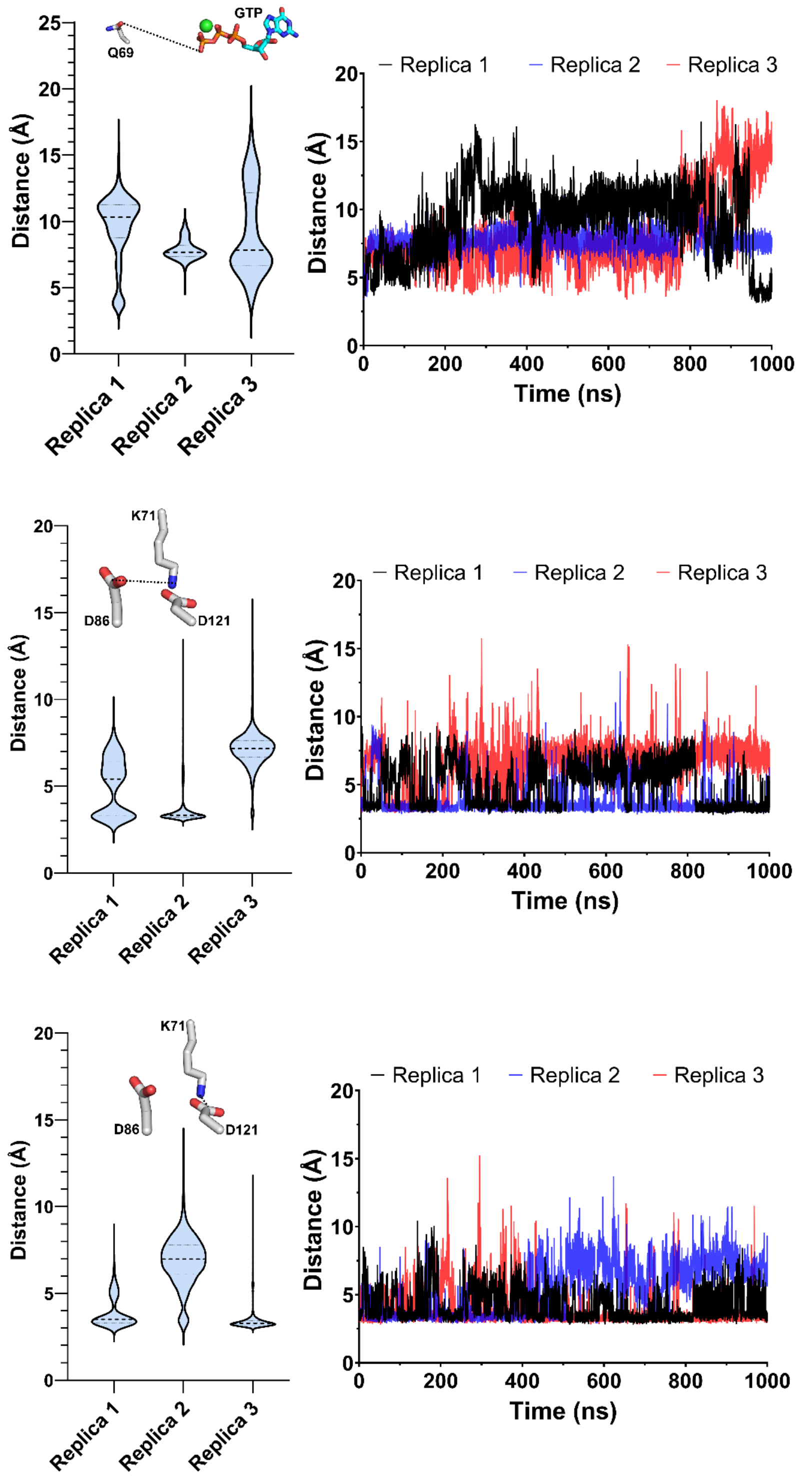


**Fig. S5. Distances between Gln69 and GTP (*top*), Asp86 and Lys71 (*middle*), and Asp121 and Lys71 (*bottom*) in the RanGAP1$^{GAP}$/Ran-GTP/RanBP2$^{RBD4}$ complex.** (*Left*) Violin plots derived from the second halves of three independent replicate simulations. (*Right*) Time evolution of the corresponding distances in each simulation.

```
  RNA1P_SCHPO ---------- ---------M SRFSIEGKSL KLDAITTEDE KSVFAVLLED DSVKEIVLSG NTIGTEAARW 51
RANGAP1_HUMAN MASEDIAKLA ETLAKTQVAG GQLSFKGKSL KLNT--AEDA KDVIKEIEDF DSLEALRLEG NTVGVEAARV 68

  RNA1P_SCHPO LSENIASKKD LEIAEFSDIF TGRVKDEIPE ALRLLLQALL K-CPKLHTVR LSDNAFGPTA QEPLIDFLSK 120
RANGAP1_HUMAN IAKALEKKSE LKRCHWSDMF TGRLRTEIPP ALISLGEGLI TAGAQLVELD LSDNAFGPDG VQGFEALLKS 138

  RNA1P_SCHPO H--TPLEHLY LHNNGLGPQA GAKIARALQE LAVNKKAKNA -PPLRSIICG RNRLENGSMK EWAKTFQSHR 187
RANGAP1_HUMAN SACFTLQELK LNNCGMGIGG GKILAAALTE CHRKSSAQGK PLALKVFVAG RNRLENDGAT ALAEAFRVIG 208

  RNA1P_SCHPO LLHTVKMVQN GIRPEGIEHL LLEGLAYCQE LKVLDLQDNT FTHLGSSALA IALKSWPNLR ELGLNDCLLS 257
RANGAP1_HUMAN TLEEVHMPQN GINHPGITAL -AQAFAVNPL LRVINLNDNT FTEKGAVAMA ETLKTLRQVE VINFGDCLVR 277

  RNA1P_SCHPO ARGAAAVVDA FSKLENIGLQ TLRLQYNEIE LDAVRTLKTV IDEKMPDLLF LELNGNRFSE EDDVVDEIRE 327
RANGAP1_HUMAN SKGAVAIADA IRGG-LPKLK ELNLSFCEIK RDAALAVAEA MAD-KAELEK LDLNGNTLGE EGC--EQLQE 343

  RNA1P_SCHPO VFSTRGRG-E LDELDDMEEL T-DEEEEDEE EEAESQSPEP ETSEEEK--E DKELADELS- -------KAH 385
RANGAP1_HUMAN VLEGFNMAKV LASLSDDEDE EEEEEGEEEE EEAEEEEEED EEEEEEEEEE EEEEPQQRGQ GEKSATPSRK 413

  RNA1P_SCHPO I--------- ---------- ---------- ---------- ---------- ---------- ---------- 386
RANGAP1_HUMAN ILDPNTGEPA PVLSSPPPAD VSTFLAFPSP EKLLRLGPKS SVLIAQQTDT SDPEKVVSAF LKVSSVFKDE 483

  RNA1P_SCHPO ---------- ---------- ---------- ---------- ---------- ---------- ---------- 386
RANGAP1_HUMAN ATVRMAVQDA VDALMQKAFN SSSFNSNTFL TRLLVHMGLL KSEDKVKAIA NLYGPLMALN HMVQQDYFPK 553

  RNA1P_SCHPO ---------- ---------- ---------- ----    386
RANGAP1_HUMAN ALAPLLLAFV TKPNSALESC SFARHSLLQT LYKV    587
```

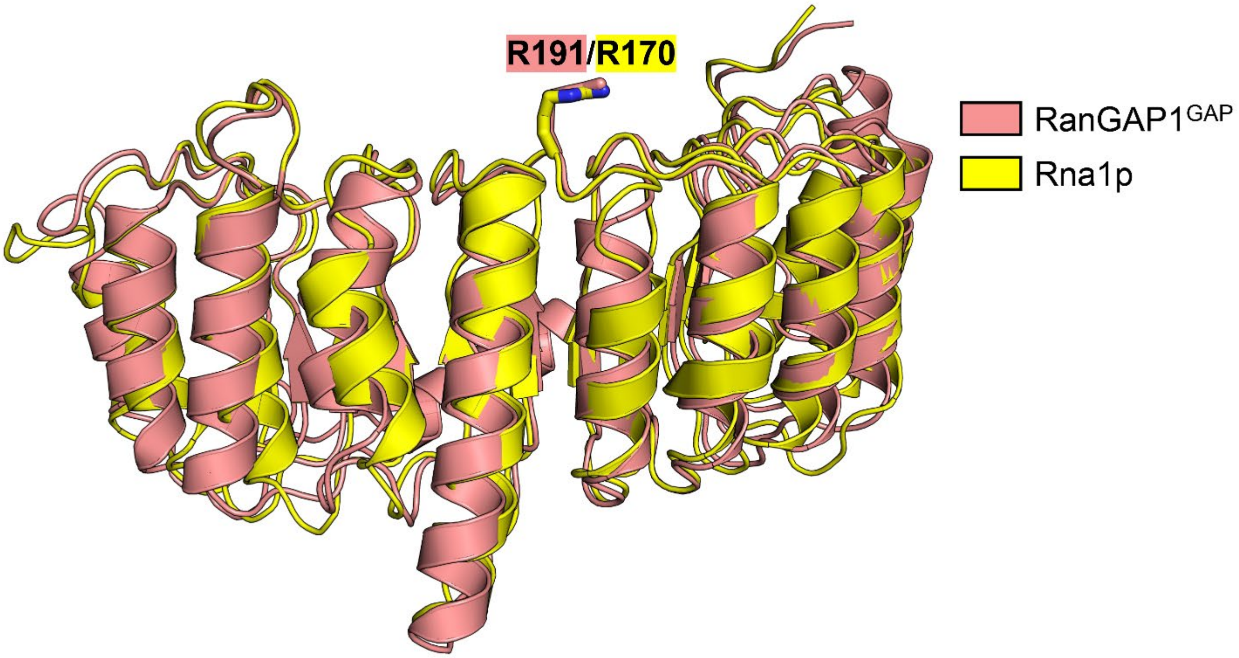


**Fig. S6. Sequence alignment of the GAP domains of RanGAP1 (human) and Rna1p (fission yeast, *Schizosaccharomyces pombe*).** Arg191 in RanGAP1 corresponds to Arg170 in Rna1p (UniProt identifier P41391). The leucine-rich repeat region of the GAP domain is highly conserved between human and yeast. The sequence identity for the GAP domains is 36%.

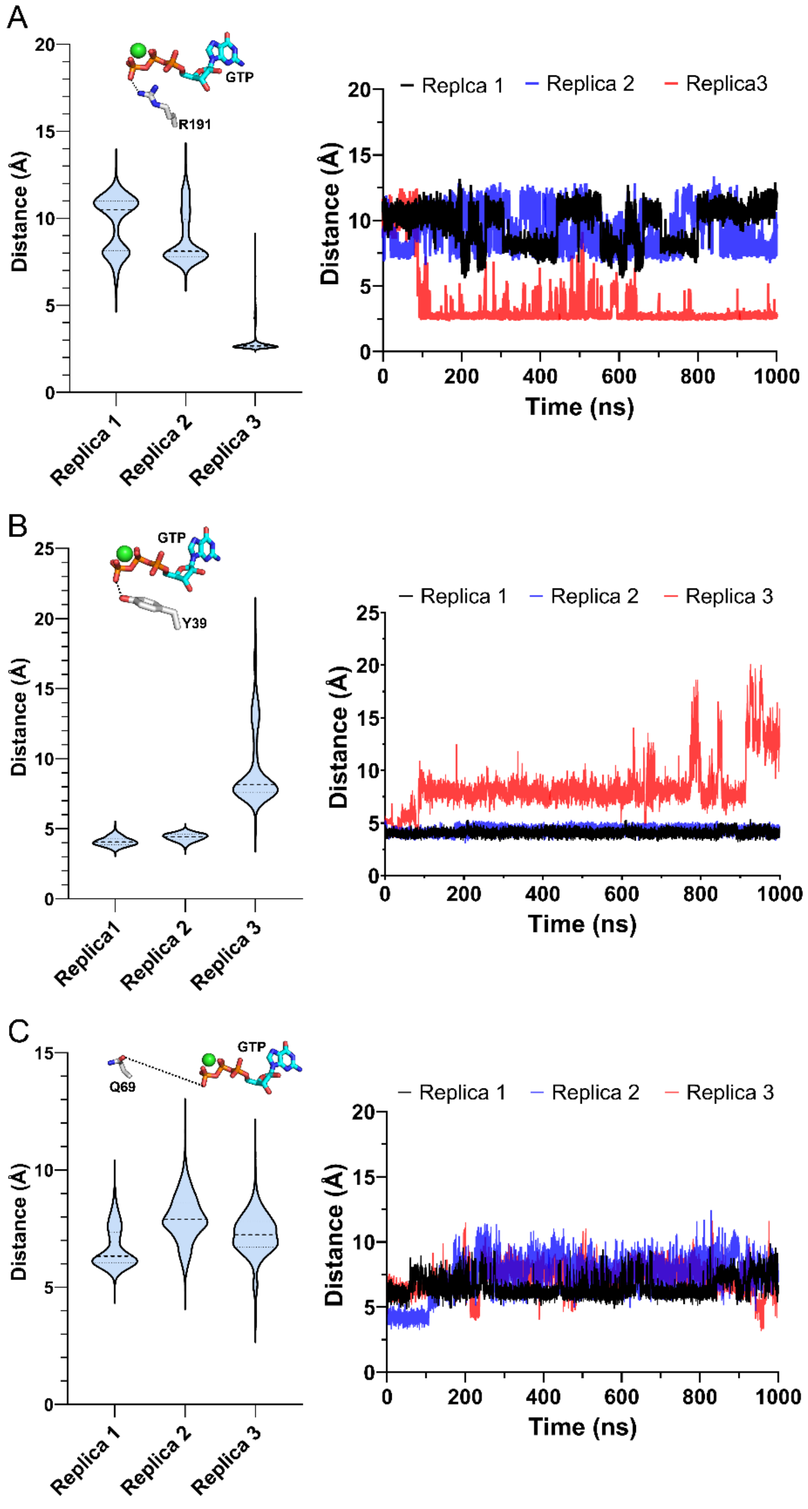


**Fig. S7. Distances between Arg191 and GTP (*top*), Tyr39 and GTP (*middle*), and Gln69 and GTP (*bottom*) in the RanGAP1$^{GAP}$/Ran-GTP/RanBP1 complex.** (*Left*) Violin plots derived from the second halves of three independent replicate simulations. (*Right*) Time evolution of the corresponding distances in each simulation.

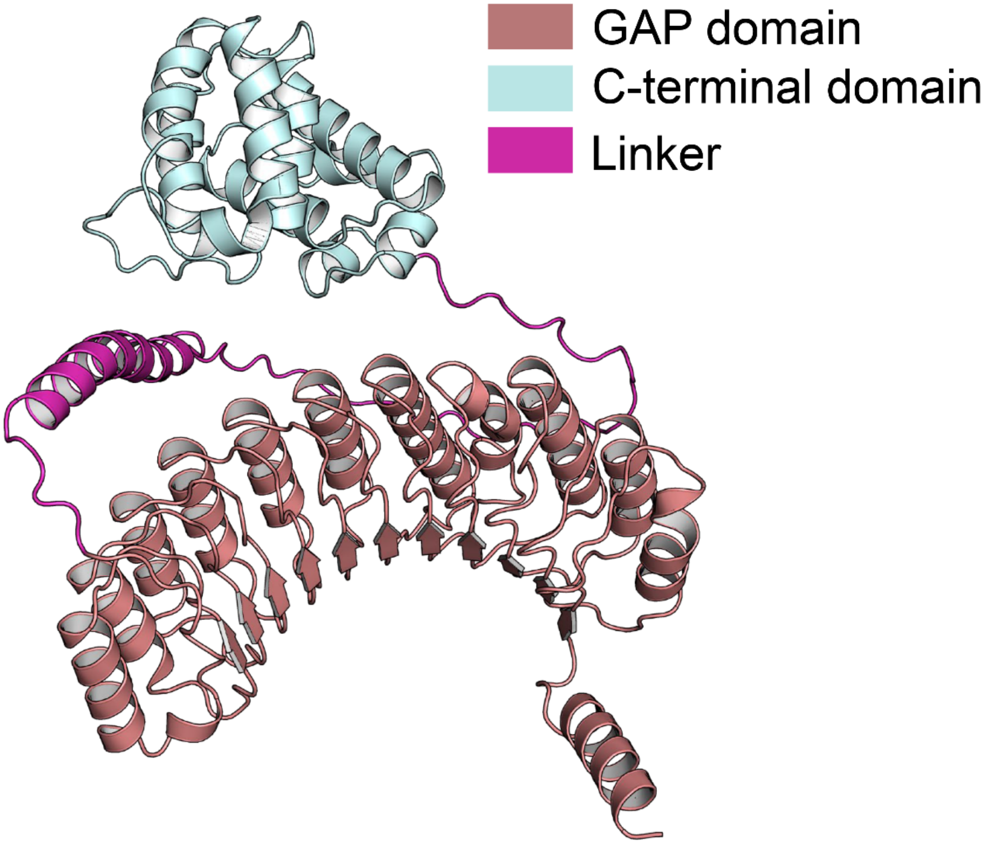


**Fig. S8. AlphaFold2 model of human RanGAP1.** No direct interactions between the GAP and C-terminal domains are observed.
